# Plant plasmodesmata bridges form through ER-dependent incomplete cytokinesis

**DOI:** 10.1101/2023.12.12.571296

**Authors:** Ziqiang P. Li, Hortense Moreau, Jules D. Petit, Tatiana Souza-Moraes, Marija Smokvarska, Jessica Perez-Sancho, Melina Petrel, Fanny Decoeur, Lysiane Brocard, Clément Chambaud, Magali Grison, Andrea Paterlini, Marie Glavier, Lucie Hoornaert, Amit S. Joshi, Etienne Gontier, William A. Prinz, Yvon Jaillais, Antoine Taly, Felix Campelo, Marie-Cécile Caillaud, Emmanuelle M. Bayer

## Abstract

Diverging from conventional cell division models, plant cells undergo incomplete division to generate plasmodesmata communication bridges between daughter cells. While fundamental for plant multicellularity, the molecular events leading to bridge stabilization, as opposed to severing, remain unknown. Using electron tomography, we mapped the transition from cell plate fenestrae to plasmodesmata. We show that the ER connects daughter cells across fenestrae, and as the cell plate matures, fenestrae contract, causing the PM to mold around constricted ER tubes. The ER’s presence prevents fenestrae fusion, forming plasmodesmata, while its absence results in closure. The ER-PM tethers MCTP3, 4, and 6 further stabilize nascent plasmodesmata during fenestrae contraction. Genetic deletion in *Arabidopsis* reduces plasmodesmata formation. Our findings reveal how plants undergo incomplete division to promote intercellular communication.

**One-Sentence Summary:** The ER is important for stabilizing nascent plasmodesmata, a process integral to incomplete cytokinesis in plants.

## Main Text

Intercellular bridges arising from incomplete cytokinesis act as structural mediators of clonal multicellularity, enabling daughter cells to communicate (*1*, *2*). These cytoplasmic connections have independently emerged across the eukaryotic tree of life spanning from animals to fungi (*3–5*). Their origin lies in the incomplete separation of daughter cells, wherein the “final cut” or abscission, responsible for severing membrane and cytosolic continuity, is impeded. Consequently, sibling cells maintain cytoplasmic bridges, forming a syncytium-like structure. While incomplete cytokinesis is cell-type specific in animals and results in a single cytoplasmic bridge, plants systematically employ this strategy to build up their communication network, creating not one but several hundreds of cytoplasmic plasmodesmata bridges, between daughter cells (Fig. 1, A to (H). These bridges are maintained post-cytokinesis and are the foundation for generating a multicellular communication network, indispensable for plant life (*6–12*).

**Fig. 1.**
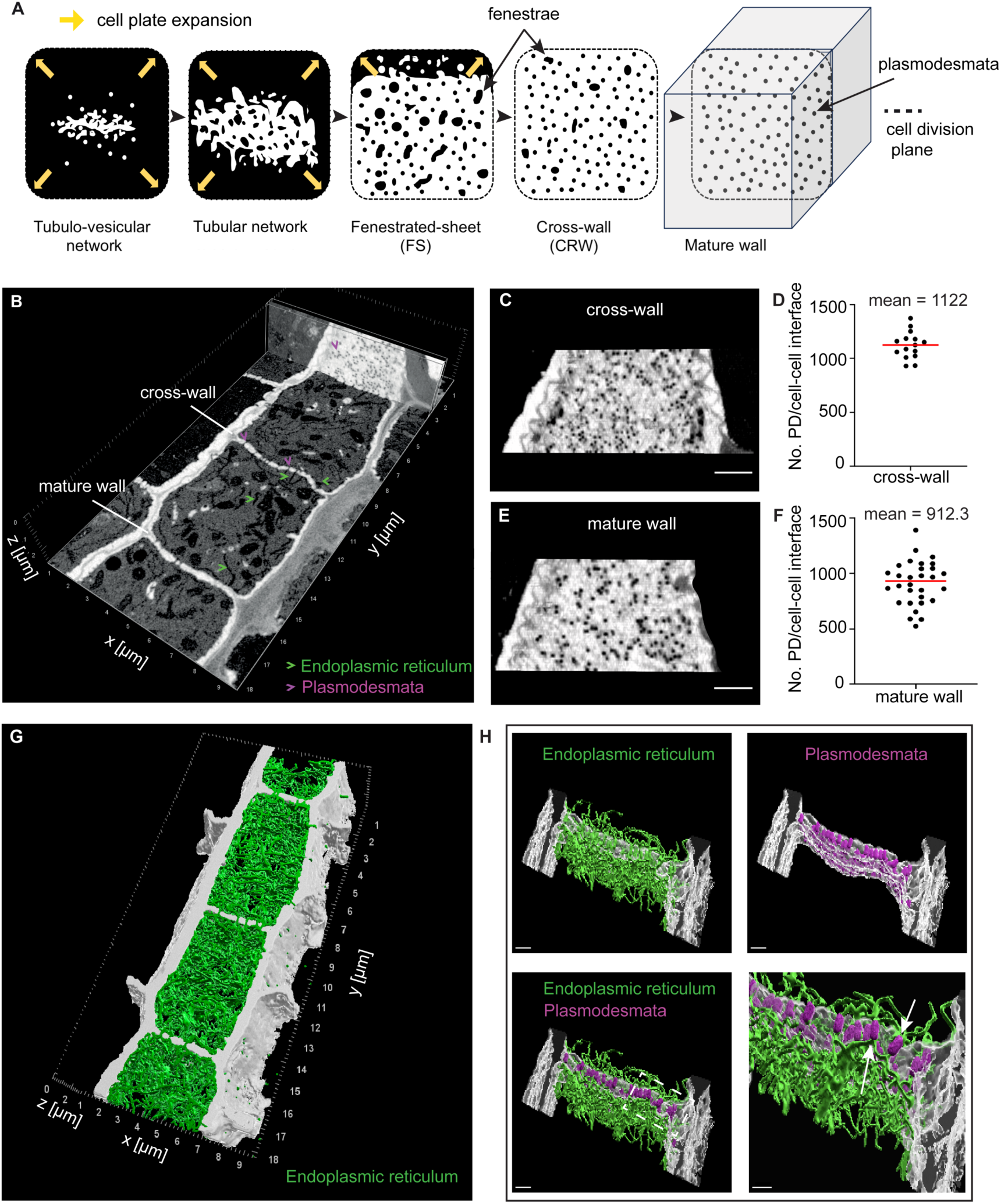
Plasmodesmata formation and ER cell-cell continuity via incomplete cytokinesis in Arabidopsis. (**A**) Schematic representation of cell plate formation during cytokinesis. (**B**) SBF-SEM image of 4-day-old *Arabidopsis thaliana* root endodermis dividing cells. (**C, E**) Orthogonal projection of a section of cross-wall (C) and mature wall (E) from B, showing plasmodesmata as black holes in the division wall (incomplete cytokinesis). (**D, F**) Plasmodesmata quantification in cross-walls (D) and mature walls (F), n = 15 and n = 30, respectively. (**G**) 3D segmentation from SBF-SEM data in (B) illustrating ER (green) continuity through adjacent cells across plasmodesmata bridges. (**H**) Zoom on cross-wall from (G) showing with ER continuity (white arrows) through plasmodesmata (magenta). Scale bar = 1 μm (C, E); 0.5 μm (three first images of panel H) and 0.4 μm (for the last image of panel H).

Vascular plants’ cytokinesis differs significantly from animals. In animals, cytokinesis involves furrowing until daughter cells remain connected by a thin intercellular plasma membrane bridge (*1*, *13*, *14*). The transition from abscission to bridge stabilization (i.e. maintaining an open cytoplasmic bridge between the daughter cells) requires ubiquitination of the ESCRT-III (endosomal sorting complex required for transport III) machinery (*15*). In vascular plant cytokinesis, a disk-shaped membrane compartment called the cell plate, expands, eventually becoming the future plasma membrane and cell wall that will separate the daughter cells (*16*, *17*) (Fig. 1A). As the cell plate forms, it contains numerous fenestrae (holes) (see fenestrated-sheet stage, Fig. 1A) that by the end of the cytokinesis (hypothetically at cross-wall stage, Fig. 1A) will either be stabilized into plasmodesmata bridges or presumably sealed off, hinting at a yet-to-be-identified molecular switch. While in animals, cytoplasmic bridge formation involves mechanisms that prevent membrane abscission between daughter cells (*15*), the molecular and cellular events underlying bridge stabilization in plants remain unexplored. We do not know how incomplete cytokinesis is achieved or how decision-making between abscission or bridge stabilization is regulated. Here, we investigated the fundamental question of how plant cells connect while dividing, using *Arabidopsis thaliana* root meristem, as an experimentally tractable model for plant cell division. Our findings reveal that the presence of the ER prevents fusion of cell plate fenestrae leading to plasmodesmata formation through incomplete cytokinesis. The multiple C2 domains and transmembrane proteins (MCTP) 3, 4, and 6 act at the ER-PM interface at contracting fenestrae, late cytokinesis, to further stabilize nascent plasmodesmata.

### Daughter cells maintain ER continuity throughout cytokinesis

An emblematic trait of plant cytokinesis lies in its capacity to preserve not just the continuity of the plasma membrane but also the continuity of the ER across division walls through plasmodesmata (*18*, *19*). Mitotic division creates a continuum of ER connections between daughter cells (Fig. 1, G and H and movie S1). Out of 126 plasmodesmata (n = 76 cross-wall, n = 50 mature wall) examined by scanning transmission electron microscopy (STEM) tomography in the division zone of the root, 120 plasmodesmata (n = 71 cross-wall, n = 49 mature wall) were confidently identified with an ER tube crossing through (fig. S1 and movie S2). While ER cell-cell continuity is a hallmark of plants, the ER’s role in incomplete cytokinesis, its dynamics, integration into plasmodesmata bridges, and its connection to bridge stability remain unclear.

With this question in mind, we first determine how the ER network becomes integrated into the forming cell wall during division. We first imaged live root meristem cells stably expressing the ER lumen (RFP-HDEL or YFP-HDEL), cell plate/plasma membrane (PM) (Lti6b-GFP) and microtubule (tagRFP-TUA5) markers (*20–22*). As the cells enter mitosis, the ER is excluded from mitotic spindles (fig. S2A) and overall, the ER pattern resembles that observed in other eukaryotes (*23*). However, starting from cytokinesis, the plant division scheme differs from animals and yeast. Membrane vesicles start gathering and fusing at the center of the division plane, and soon form a disk-shaped membrane compartment punctured by fenestrae. The cell plate then expand centrifugally to finally partition the daughter cells (*16*). By simultaneously tracing cell plate vesicles (Lti6b-GFP) and the ER (RFP-HDEL), we observed early ER accumulation at the cell plate (fig. S2B and movie S3). High-resolution airyscan imaging revealed ER strands crossing the cell plate throughout cytokinesis (fig. S3, A and B), resembling the dense ER intercellular matrix connecting post-cytokinetic cells (Fig. 1 G and H, fig. S1, and movies S1 and S2). At early stages of cytokinesis, the ER was already one single and continuous compartment, stretching across the two daughter cells as demonstrated by fluorescence loss in photobleaching (FLIP) targeting the ER luminal marker (YFP-HDEL) (fig. S3, C and D and movie S4). Thus, cell-cell ER continuity originates from early cytokinesis. The ER is anchored in both daughter cells, bridging them across the cell plate.

### Plasmodesmata stabilize in the presence of the ER

We then looked at the fate of the fenestrae in relation to the ER. Previous work showed ER association with fenestrae, from which plasmodesmata were proposed to originate (*17*). To understand the relationship between the ER and fenestrae, we examined the fate of each in relation to the other. For that, we looked back at fenestrae events along the entire division plane using electron tomography (movie S5). We focus on both fenestrated-sheet stage, when the cell plate consists of an almost continuous membrane not yet fused to the parental walls, and the cross-wall stage, when the cell plate has completed fusion, and fenestrae are transitioned into the plasmodesmata (Fig. 1A and Fig. 2A; fig. S4). We employed chemically fixed and osmium ferricyanide-stained root meristem to specifically enhance ER staining and clearly visualize this membrane compartment during cell plate formation (*19*). In total, we observed 118 fenestrae events by electron tomography from five and four cytokinetic cells for fenestrated-sheet and cross-wall stages respectively, providing a comprehensive and quantitative representation of plasmodesmata formation (Fig. 2 and Fig. 3). We found a marked disparity between the two stages regarding ER association with fenestrae. At cross-wall stage, all fenestrae presented continuous ER. In contrast, in preceding fenestrated-sheet stage, only 65.7% of fenestrae showed clear ER physical continuity, 11.8% had ER association without cell-cell continuity and the remaining 22.5% showed no ER continuity with fenestrae sealing off (Fig. 2, B and C, and movies S6 to S8). Thus, not all fenestrae exhibit ER association, but all plasmodesmata contain ER.

**Fig. 2.**
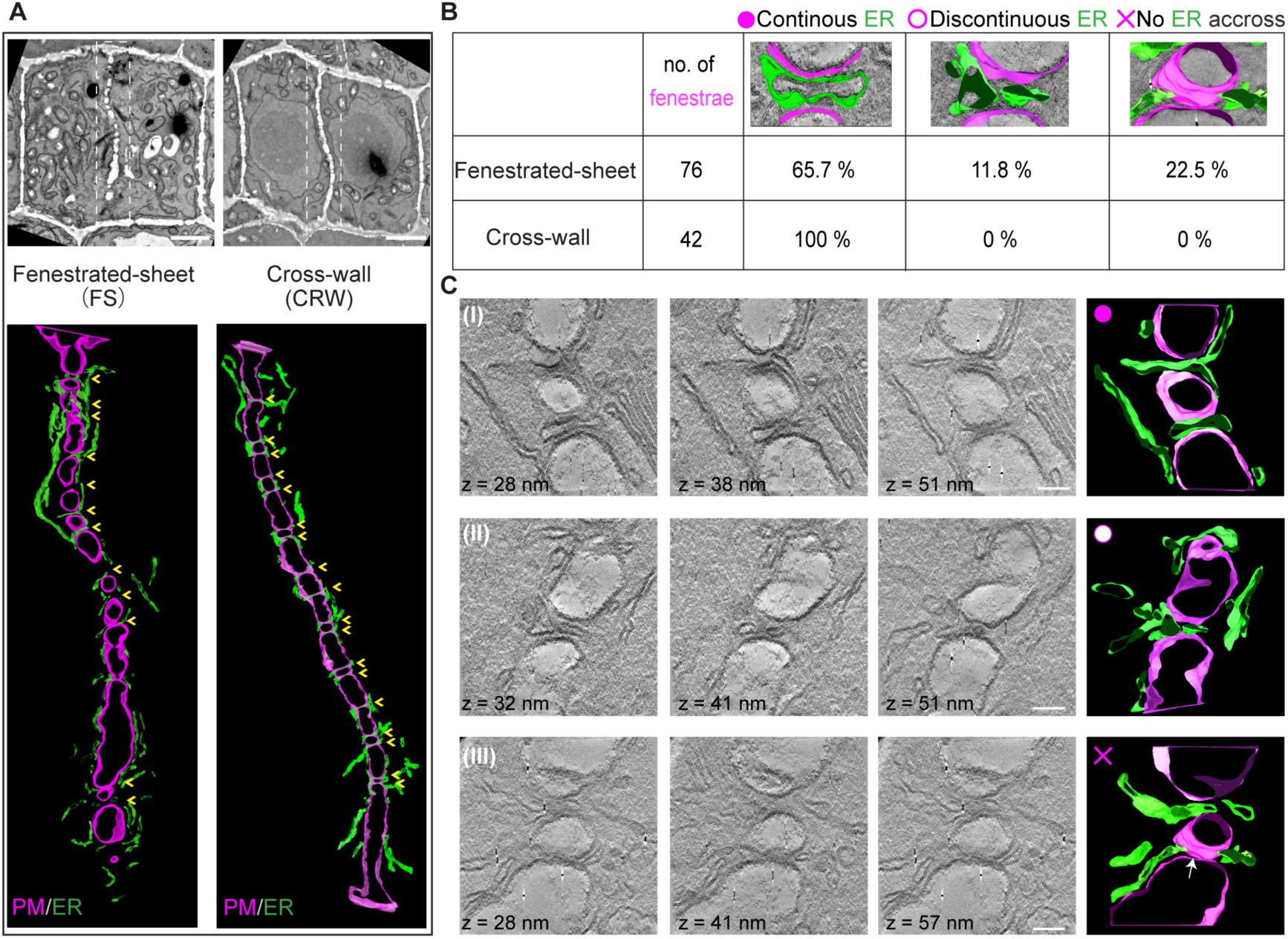
Ultrastructural observations of forming plasmodesmata. (**A**) Batch electron tomography acquisitions along the entire cell plate at fenestrated-sheet (FS, n = 5) and cross-wall (CRW, n = 4) stages (top, overview of the dividing cells; bottom 3D segmentation of the cell plate for stitched tomograms). Cell plate membrane (PM) in magenta, ER in green, yellow arrows point to fenestrae. (**B**) Quantification of fenestrae events presenting continuous ER across, discontinuous ER across or no ER across at fenestrated-sheet and cross-wall stage. n = 76 fenestrae for FS (five complete cell plates) and n = 42 fenestrae for CRW (four complete cell plates). (**C**) Reconstructed tomography sections across fenestrae events and 3D segmentation showing: (I) open fenestrae with continuous ER; (II) open fenestrae with discontinuous ER and (III) closing fenestrae with ER structures flanking the fusion site (white arrow). Scale bars, 2 μm (A); 100 nm (C).

**Fig. 3.**
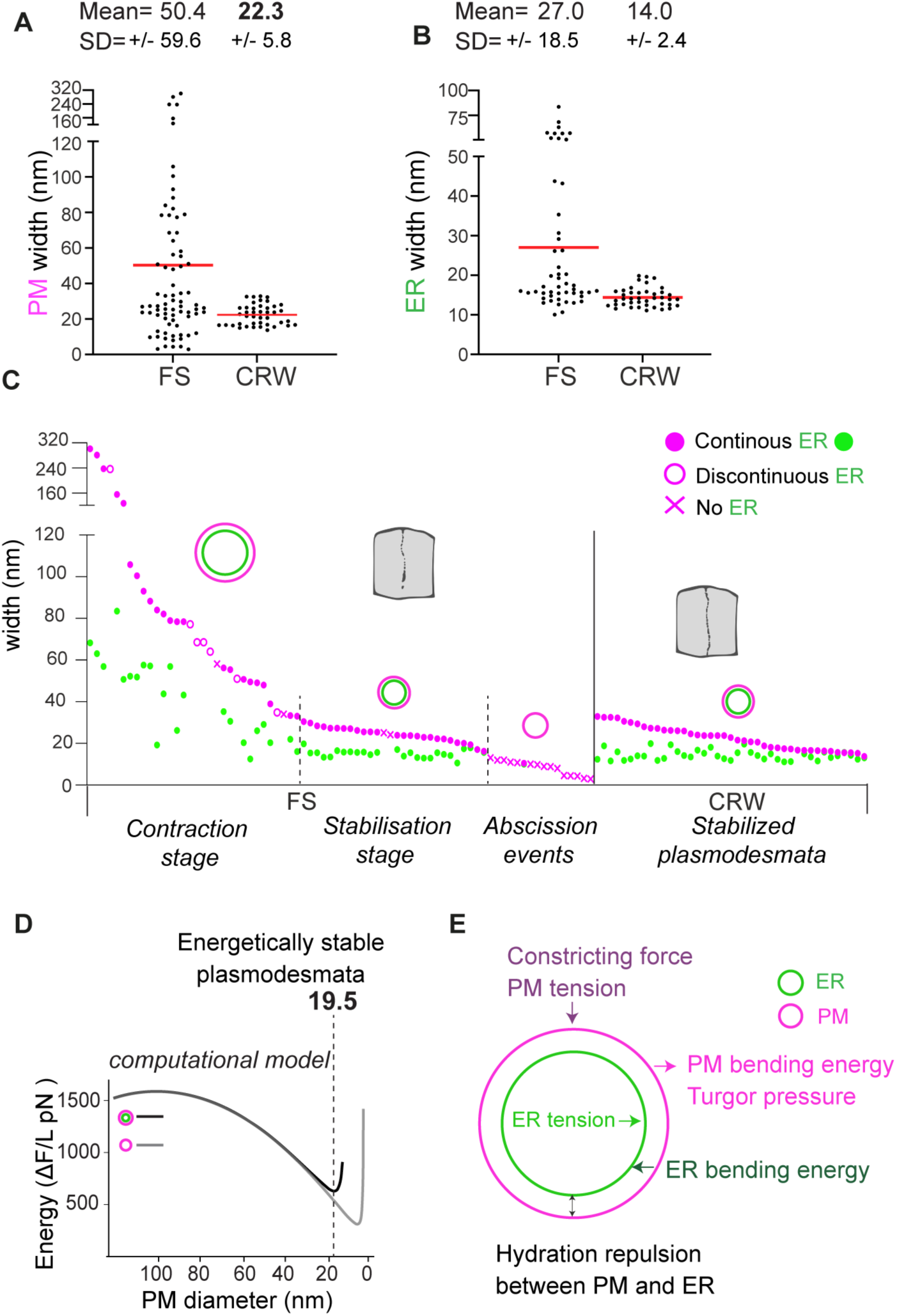
Fate of the fenestrae in relation to the ER. (**A**) Diameter of fenestrae (PM) at fenestrated-sheet (FS) and cross-wall (CRW) stages, n = 76 for FS and n = 42 for CRW extracted from electron tomography acquisition from Fig.2 (n= 5 (FS stage) and n = 4 (CRW stage) cells). (**B**) Diameter of ER tubes across fenestrae at FS and CRW stages. n = 50 for FS and n = 41 for CRW extracted from electron tomography acquisition from Fig.2. (**C**) Plotting of fenestrae (PM) diameters together with ER state (continuous, discontinuous, absence, and diameter) during FS and CRW stages, PM in magenta and ER in green. (**D**) Computational model of energetically metastable plasmodesmata. The presence of ER works against full fenestrae closure (PM sealing) by creating an extra energy barrier (that of ER fission or removal), leading to metastable structure of about 19.5 nm in diameter (ΔF, free energy of the structure. L, longitudinal length of the fenestrae - see suppl. text for detailed explanations). (**E**) Schematic representation of the various terms taken into consideration to model the free energy of the system with the fenestrae in magenta and the ER in green.

### Fenestrae close around the ER, forming uniform-dimension plasmodesmata

During cell plate maturation, fenestrae are thought to shrink until they formed plasmodesmata or are sealed off. (*17*). This implies that one dividing cell can engage in both abscission and stabilization events while it progresses to the end of cytokinesis. To provide more details of the stages of plasmodesmata maturation, we systematically correlated the size of fenestrae with the ER in its various states (continuous, discontinuous, no ER) across both fenestrated-sheet and cross-wall stages (Fig.3, A to C). We indeed observed contraction of fenestrae as the cell plate matures (Fig. 3, A to C). At fenestrated-sheet stage, fenestrae diameter spans from 301.2 nm to below 10 nm. By the end of cytokinesis (cross-wall stage), fenestrae had stabilized to a uniform diameter of 22.3 ± 5.8 nm (mean ± SD) (Fig. 3A, and fig. S1). The stabilization of contracting fenestrae into plasmodesmata bridges was invariably associated with the presence of ER in a contracted form (Fig. 3B, and fig. S1). In the absence of the ER, fenestrae diameter decreases below 20 nm until complete closure (as they are not present in cross-wall stage) (Fig. 3C). Thus, during cell plate maturation, fenestrae constrict and mold around the ER, leading to plasmodesmata. Fenestrae with no ER are not maintained.

### Modeling fenestrae closure in relation to the ER

Our data suggest a regulated process where fenestrae can only be stabilized into plasmodesmata (i.e., not be sealed off) in the presence of the ER. To understand the plausible physical-basis behind this process, we built a semi-quantitative physical and computational model of plasmodesmata formation. Our model computes the free energy of the system (Fig. 3D), in our case a single bridge, and includes the contributions of *i)* the cell plate (membrane bending energy and lateral tension, turgor pressure, and a force associated to cell plate expansion), *ii)* the ER (tubule bending energy and lateral tension); and *iii*) an interaction term due to hydration repulsion between the ER and cell plate membranes (see Fig. 3E, fig. S5 and suppl. text for a detailed discussion).

The model outputs suggest that the cell plate’s expansion initially energetically favors fenestrae shrinking, up to a point (minimum of the free energy profile; Fig. 3D) when the membranes are getting close to each other, and extreme bending and hydration-repulsion energy dominate, stopping fenestrae shrinking. From this point, to seal fenestrae the system needs to overcome two barriers; one associated with ER removal (tube removal/fission) and a second one associated with fenestrae sealing. Moreover, the hydration-repulsion between the fenestrae and ER membranes also works against the fusion between of these two membranes. Altogether this results in fenestrae constriction being stopped at larger pore sizes compared to what would happen without ER (7 nm without ER to 19.5 nm with ER; Fig. 3D). Therefore, in the presence of the ER, fenestrae first reach an energetically metastable state (which does not exist without the ER) with a predicted diameter of 19.5 nm (Fig. 3D), closely matching experimental data (22.3 nm, Fig. 3A). In this view, the ER helps prevent full fenestrae closure, as the energy required to break the ER tube is not provided. This causes the PM to mold around the ER to achieve the lowest energy state. This could explain why plasmodesmata consistently include ER and maintain a uniform diameter.

### MCTP3, 4, 6 ER-tethers contribute to plasmodesmata formation

Given the ER’s dynamic nature, specific factors are likely required to stabilize the ER while plasmodesmata form. Previously, we identified, a plasmodesmata-specific ER protein family, MCTPs, that function as ER-PM tethers (*24*). MCTPs are among the few plasmodesmata-enriched ER proteins identified so far and were proposed to be core structural elements (*24*, *25*). According to single-cell RNA sequencing (*26*), among 16 Arabidopsis members, MCTP3, MCTP4, MCTP6, and MCTP7 were expressed in dividing root meristematic cells (fig. S6A). When fluorescently-tagged and expressed under endogenous promoters, only MCTP3, MCTP4, and MCTP6 exhibited broad root expression and localized to the ER/cell plate in dividing cells (fig. S6, B to E). They also appeared as dots, typical of plasmodesmata association, at cell-cell interfaces of newborn daughter cells which we confirmed by correlative light and electron microscopy (CLEM) for MCTP4 (Fig. 4, A and B). High-resolution airyscan microscopy, revealed that the ’dotty’ MCTP-fluorescent signals were actually stripes extending across the wall connecting sister cells (Fig. 4C). In contrast, proteins like synaptotagmin (SYT1 and SYT5) and reticulon (RTN 6), which are involved in ER-PM tethering and shaping (*27*, *28*), were absent from the plasmodesmata (Fig. 4C). These findings underscore the molecular specialization of ER within newborn cytokinetic-plasmodesmata.

**Fig. 4.**
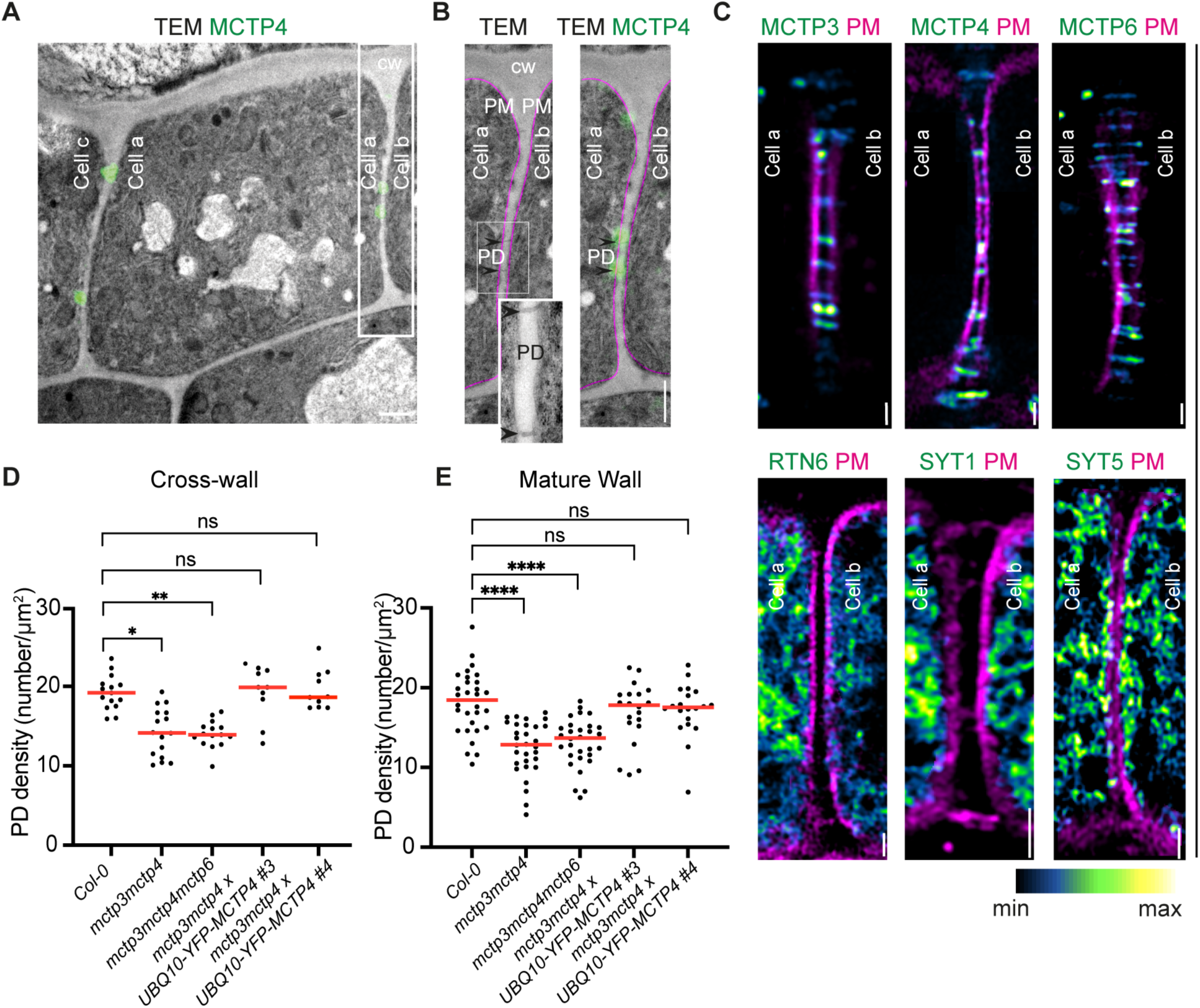
MCTPs are important factors of plasmodesmata formation. (**A-B**) CLEM on YFP-MCTP4 expressed in genetically complemented *mctp3mctp4* mutants in root meristem epidermal cells. (B) Enlarged region as in A (outlined white rectangle). Black arrows point to the two plasmodesmata connecting cell a and cell b. Green: YFP-MCTP4; Magenta: PM; CW: cell wall; TEM: transmission electron microscopy. (**C**) airyscan imaging of live Arabidopsis meristem epidermal cells expressing YFP-tagged MCTP3, MCTP4, MCTP6, SYT1, STY5 (green fire blue) under their native promoter; RTN6 under 35S promoter. PM (magenta) is stained by FM4-64. (**D-E**) Quantification of plasmodesmata density on cross-walls and mature walls (number of plasmodesmata/μm^2^) in the *A. thaliana* root meristem endodermis cells using SBF-SEM. The bars indicate the mean. Significance was tested using ordinary two tailed Mann-Whitney U-tests (****, P<0.0001). n = 15 (Col-0), n = 16 *(mctp3mctp4*), n = 15 (*mctp3mctp4mctp6*), n = 10 (*mctp3mctp4 x UBQ10-YFP-MCTP4 line* #3), n = 10 (*mctp3mctp4 x UBQ10-YFP-MCTP4 line #4*) cells for the cross-wall quantification. n = 30 (Col-0), n = 30 (*mctp3mctp4*), n = 30 (*mctp3mctp4mctp6*), n = 20 (*mctp3mctp4 x UBQ10-YFP-MCTP4* line #3), n = 20 (*mctp3mctp4 x UBQ10-YFP-MCTP4* line #4) cells for the mature wall quantification. Scale bars, 1 μm (A, B, C).

To test if ER-PM contact is indeed playing a role in bridge stabilization, we generated *mctp* loss-of-function mutants and asked whether they present a defect in plasmodesmata production. Previous work reported a general growth and development defect of *mctp3mctp4* mutant consistent with a hypothetical defect in plasmodesmata formation (*24*). As MCTP3 and MCTP4 share 98.7% similarity in amino acids and are functionally redundant (*24*), we focused on the double (*mctp3mctp4*) and triple (*mctp3mctp4mctp6*) higher-order mutants. We first quantified plasmodesmata in newly formed post-cytokinetic walls across four root cell layers (epidermis, cortex, endodermis and pericycle - apico-basal post-division walls) using transmission electron microscopy. We found that *mctp* double and triple mutants presented a significant drop of plasmodesmata in all four layers examined (reflected as density, number per unit area, ranging from 28% drop in the cortex to 47% drop in the epidermis) (fig. S7) when compared to the wild-type. The reduced but not complete absence of plasmodesmata could be attributed to redundancy, given the extensive MCTP multigenic family and potential involvement of other factors. While we speculated that the plasmodesmata phenotype in the *mctp* higher-order mutants originate from cytokinesis defects, selective removal of mature post-cytokinetic plasmodesmata cannot be ruled out. We reason that cytokinesis-defect should reduce both cross-wall and mature wall plasmodesmata, while post-cytokinesis elimination would impact mature walls only. To test that, we took advantage of serial block-face scanning electron microscopy (SBF-SEM) to map plasmodesmata across the entire cell volume at cross-wall and mature wall stages (fig. S8). We focused on the endodermis due to its superior sample preservation. SBF-SEM data indicate that the plasmodesmata deficiency originated from cytokinesis as *mctp* double and triple mutants presented significantly fewer cytoplasmic bridges from cross-wall stage (Fig. 4, D and E). Complementation of the double mutant with a *UBQ10:YFP-MCTP4* transgene was sufficient to restore plasmodesmata to the wild-type level (two independent lines; Fig. 4, D and E). No general cytokinesis defects, such as aborted cell plate or branching or misalignment of the cell plate, were observed in *mctp* mutants nor did we observe obvious defect in ER accumulation at the cell plate during cytokinesis (fig. S9). Our results show that ER-associated MCTP tethers are needed for efficient plasmodesmata formation during cytokinesis.

### MCTP3,4,6 further stabilize contracting ER-containing fenestrae

Next, we enquired how MCTPs may contribute to plasmodesmata formation. Using live imaging, we first followed MCTP dynamics during cytokinesis in genetically complemented plants. MCTP3, 4 and 6 proteins initially showed a uniform distribution at the ER-associated cell plate, comparable to the general ER membrane marker cinnamate 4-hydroxylase C4H-GFP (*29*) (Fig. 5A, fig. S10, A to C). However, 5 to 10 min before cross-wall transition (annotated as t=0), YFP-MCTPs signal started to cluster as dots, contrasting with C4H-GFP’s uniform distribution. These YFP-MCTP’s dots correspond to nascent plasmodesmata, as confirmed by CLEM (Fig. 5B; six plasmodesmata out of two cross-walls). MCTP clustering aligns with fenestrae stabilization into plasmodesmata, when the PM wraps around a constricted ER tube (Fig. 3; fig. S1), indicating a link between ER-cell plate tethering and ER constriction.

**Fig. 5.**
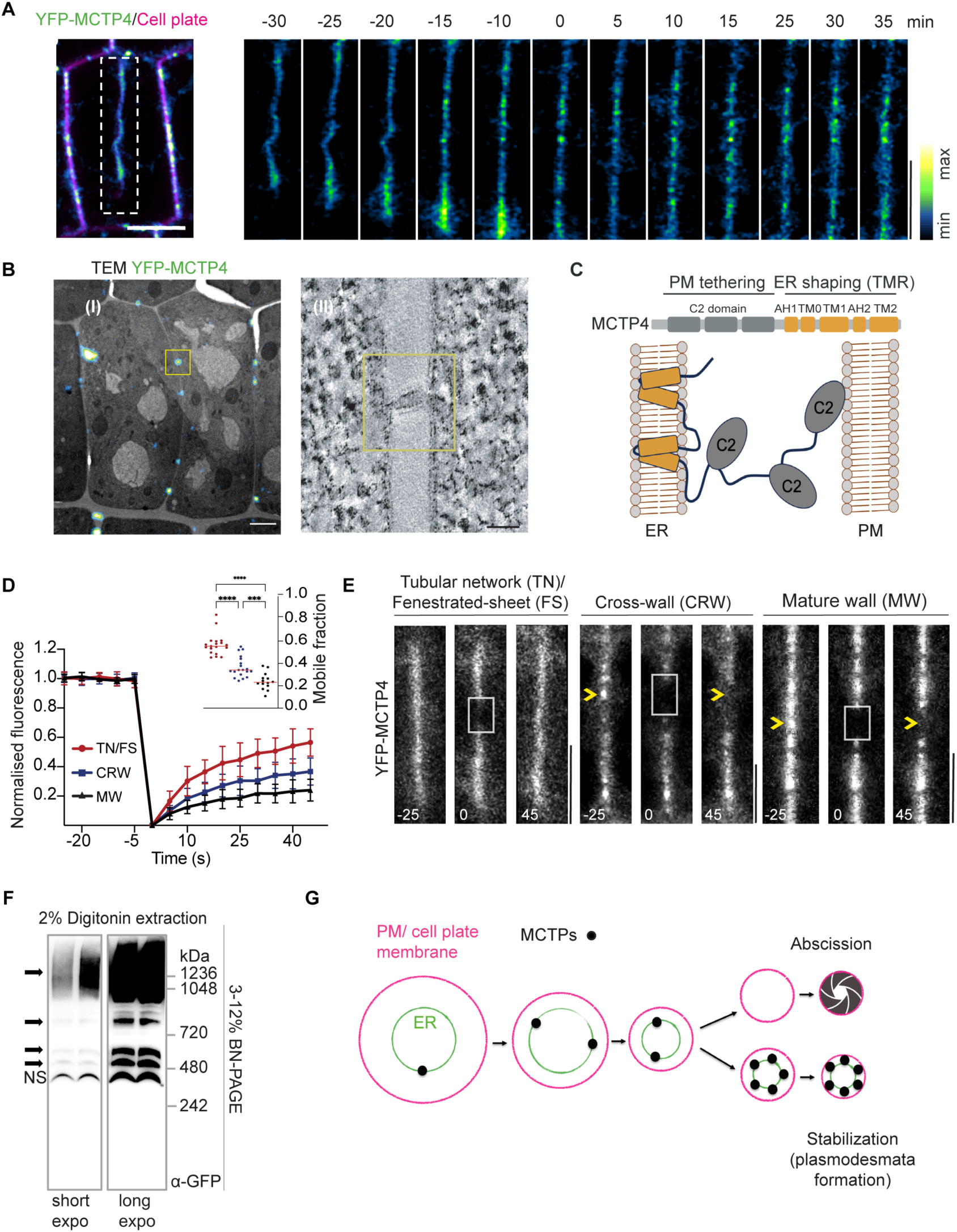
MCTPs cluster within nascent plasmodesmata and aid stabilization. (**A**) YFP-MCTP4 dynamics during cytokinesis in *mctp3mctp4* complemented lines root epidermis. (CRW stage = time 0). Cell plate is labelled by FM4-64. (**B**) Analyzed by CLEM, YFP-MCTP clusters (I) correspond to plasmodesmata (II). (**C**) Schematic illustration of MCTP domains. (**D-E**) YFP-MCTP4 mobility, measured by FRAP. The white squares indicate the photobleached regions. Yellow arrows indicate MCTP4 ’dots’ that do not recover fluorescent signal. Quantification of fluorescence (mean ± SD) and mobile fraction (bars indicate mean) in tubular network (TN)/FS (n = 19), CRW (n = 19) and MW (n = 14) stages (****, P<0.0001; two tailed Mann-Whitney U-tests). (**F**) Blue native-PAGE of YFP-MCTP4. Black arrows show MCTP4 high-molecular-weight complexes in short (51 s) and long exposed (157 s) blots. Non-specific (NS) indicates non-specific band (N = 3). (**G**) Conceptual model of ER and MCTP action during plasmodesmata formation. Scale bars, 5 μm (A and E), 2 μm (B I), and 100 nm (B II).

We therefore investigated if, in addition to their membrane tethering activity, MCTPs could also shape the ER. We found that the ER-anchor C-terminal region of MCTPs (transmembrane region, TMR), presents homology to the ER-shaping reticulon-domain (*30*), a function we experimentally confirmed *in planta* and in yeast (Fig. 5C, fig. S11). To examine the role of MCTP ER-shaping and C2 lipid-binding domains in targeting plasmodesmata, we generated truncated mutants with sequential deletions (fig. S12A). Except for the C2B deletion mutant, clustering at plasmodesmata was lost in all tested truncations (fig. S12, B and C), indicating that both the TMR domain (ER shaping) and C2 cytoplasmic domain (membrane tethering) are important for accumulation at nascent plasmodesmata.

The molecular machinery for abscission/stabilization in animals and yeast operates locally within the bridge, adopting a stationary matrix-like molecular organization (*13*, *31*). We speculated that MCTPs behave alike and investigated their mobility by fluorescence recovery after photobleaching (FRAP) and FLIP. These techniques complement each other, with FRAP showing the mobile protein fraction and its dynamics, while FLIP reveals the locations of non-mobile proteins by depleting the mobile population. All three MCTPs exhibited high-dynamic behavior before accumulating at nascent plasmodesmata (Fig. 5, D and E and fig. S13, A to C). However, from the cross-wall stage, their mobility dramatically decreased as indicated by FRAP, and in nascent-plasmodesmata associated MCTP signal persists under FLIP (fig. S13, D and E). In contrast, the ER membrane marker, C4H-GFP, remained highly mobile throughout cytokinesis (fig. S13A).

A straightforward explanation for the stable accumulation of MCTPs at contracting fenestrae is their organization in polymeric lattices. Such hypothesis is consistent with the presence of an RTN-homology domain known to induce oligomerization (*32*). Using co-immunoprecipitation, we show that all three MCTP RTN-homology domains can indeed physically interact (fig. S14A). To further test whether MCTPs form oligomeric complexes, we extracted YFP-MCTP4 (from *A. thaliana mctp3mctp4* complemented lines) from plasmodesmata-enriched wall fractions under native extraction conditions, using non-denaturing native-blue gel. The majority of YFP-MCTP4 (120kD as monomer) was detected as a complex of about 1000 kDa regardless of the non-ionic detergents used (Fig 5F and fig. S14, B to C). Strikingly, YFP-MCTP4 high-molecular-weight complexes resisted strong disruptive SDS/DTT solubilization conditions (fig. S14, D and E). Thus, plasmodesmata-enriched MCTPs form highly stable oligomer complexes.

Based on our observations, we hypothesized that ER-associated MCTPs concentrate at fenestrae ingression sites, where they oligomerize, stabilizing ER strands across nascent bridges and forming a protective shield to prevent abscission. In this scenario, loss of MCTPs would shift the balance towards abscission, explaining the loss of plasmodesmata (Fig. 4, D and E, and fig. S7). To assess the feasibility of this hypothesis, we updated our semi-quantitative physical model by incorporating MCTPs (see Suppl. text for details). The model predicts that the enrichment of MCTPs in contracting fenestrae is energetically favorable, as the system free energy decreases with increasing concentration of functional MCTPs (i.e., with ER-shaping and/or membrane tethering function) in the membrane bridge (fig. S15, B). This sorting effect may arise from both the curvature-generating/sensing characteristics of the RTN-homology domain and the contact-driven sorting through tethering activity. According to our physical model, the presence of an ER tubule, within narrowing fenestrae, creates an energy barrier (that of ER tubule fission or ER-tubule removal) working against fenestrae closure (Fig. 3D). MCTPs introduce an extra obstacle compared to the ER alone, further hindering complete closure and leading to the formation of metastable ER-cell plate MCTP-rich membrane bridges. The model further predicts that stabilized plasmodesmata should display the same diameter with or without MCTPs (fig. S15, C). To check this prediction, we measured nascent plasmodesmata diameter at cross-wall stage (when they are stabilized) in mctp3mctp4mctp6. The *mctp* mutant displays an average diameter of 22.3 nm ± 5.7 nm (mean ± SD) (fig. S16) similar to the wild-type (Fig. 3A). Collectively, our data support a spatio-temporal coordination model, wherein MCTPs concentrate at fenestrae ingression sites to assist the ER’s action and establish stable communication bridges (Fig. 5G).

## Conclusion

In this study, we identified a novel function of the ER in incomplete cytokinesis in plants, and plasmodesmata bridge formation. Abscission and stabilization events occur concurrently during the same cytokinetic event, and the switch between the two is regulated by the ER along with MCTP proteins (Fig. 5G). Our observations resolve the longstanding puzzle regarding the presence of ER inside plasmodesmata and highlights the necessity for intercellular ER continuity.

## Acknowledgments

We would like to thank Kevin Verstaen from the Vlaams Instituut voor Biotechnologie single cell core who provided the single cell sequencing raw data, which we used in the fig. S6. Sofia Otero provided the MCTP6 native reporter line. We thank Sebastian Marais for the help with iMaris software. Guillaume Maucort for the suggestions on the SBF-SEM visualization. Fabrice Cordelières for live-imaging data display. We would also like to thank Maya Schuldiner, Patricia Bassereau, Olivier Hamant, Agathe Chaigne, Thibaut Brunet, Yohann Boutté, Sébastien Mongrand for reading and commenting on the article. All light and electron imaging were done at the Bordeaux Imaging Center, member of the national infrastructure France-BioImaging supported by the French National Research Agency (ANR-10-INBS-04).

## Author contributions

Conceptualization: ZPL, EMB

Methodology: ZPL, HM, JDP, MP, LB, EG, FC, EMB

Validation: ZPL, HM, JDP, EMB

Investigation: ZPL, HM, JDP, TSM, MS, JPS, MP, FD, LB, CC, MG, AP, MG, LH, ASJ, EG, FC

Visualization: ZPL, HM, JDP, FC, EMB

Supervision: EMB

Funding acquisition: MCC, EMB

Writing – original draft: ZPL, EMB

Writing – review & editing: ZPL, HM, FC, YJ, AT, WAP, MCC, EMB

## Data and materials availability

All data are available in the manuscript or the supplementary material and all materials are available upon request. Wolfram mathematice notebook for the physical modeling is available on GitHub (citation) as well as Zenodo repository (citation).

F. Campelo, Wolfram mathematice notebook used to compute the results of the plasmodesmata model in Li, Moreau et al., 2024.https://github.com/ziqiangpatrickli/fenestraephysicalmodeling.git

F. Campelo, Wolfram mathematice notebook used to compute the results of the plasmodesmata model in Li, Moreau et al., 2024. https://zenodo.org/records/12746245

## Supplementary Materials

### Materials and Methods

#### Plasmid constructs

Constructs used in this study including MCTP3 (At3g57880), MCTP4 (At1g51570), MCTP6 (At1g22620), and MCTP7 (At4g11610) were cloned from Arabidopsis Col-0 cDNA/genomic or synthesized DNA. All promoters were cloned into pDONR-P4RP1; genes and fluorescent tags were cloned into either pDONR221 or pDONR-P2RP3 using the GATEWAY cloning system. Domain deletion mutants were generated by Gibson assembly and introduced in the entry and destination constructs using Gateway assembly. All constructs were assembled using MultiSite-Gateway reactions where three segments were cloned into the destination vector, pLOK180, which provides a red seed-coat selection marker (FAST-red cassette) (*35*).

#### Plant material and growth conditions

All the *Arabidopsis thaliana* transgenic lines used in this study were generated from the Columbia-0 accession. *mctp3* (Sail-755-G08) and *mctp4* (Salk-089046) single T-DNA insertion mutant and *mctp3mctp4* double mutant were described in (*24*). The triple mutant *mctp3mctp4mctp6* results from a cross between *mctp3mctp4* and *mctp4mctp6,* the latter was generated by crossing *mctp4* and *mctp6* single mutant (SALK_145386C). GV3101 agrobacterium strains expressing the constructs of interest were used to transform *Arabidopsis* Col-0 and *mctp* mutants by floral dip (*36*). Transformed seeds were selected under an epi-illumination Axiozoom microscope (Zeiss) based on the red seed coat selection (*35*). Arabidopsis lines were grown vertically at 22°C in long days light conditions (16-h light/8-h dark cycle with 70% relative humidity and a light intensity of 200 µmol. m^-2^. s^-1)^ on solid half-strength Murashige and Skoog (½ MS) supplemented with vitamins (2.15 g/L), MES (0.5 g/L), sucrose (10 g/L) and plant agar (7g/L), pH 5.7. *Nicotiana benthamiana* plants were cultivated in the greenhouse (18/25°C night/day).

#### Transient expression in *Nicotiana benthamiana*

For transient expression in *N. benthamiana*, leaves of 3 to 4-week-old plants were pressure-infiltrated with GV3101 agrobacterium strains, previously electroporated with the relevant binary plasmids. Before infiltration, agrobacteria cultures were grown in Luria and Bertani medium with appropriate antibiotics at 28°C for 2 days, then diluted to 1/10 and grown until the culture reached an OD600 of about 0.6-0.8. Bacteria were then pelleted and resuspended in water at a final OD600 of 0.3 for individual constructs and 0.2 each for the combination of the two. The ectopic silencing suppressor P19 with an OD600 of 0.05 was co-infiltrated. Agroinfiltrated N. *benthamiana* leaves were imaged 2–3 days post-infiltration at room temperature.

#### Bacterial strains

Escherichia coli DH5-Alpha, Stbl2, and Agrobacterium tumefaciens GV3101 strains were from laboratory stocks.

#### Transmission electron microscopy (TEM) sample preparation, imaging and plasmodesmata quantification

4-day-old Arabidopsis seedling roots were chemically fixed at RT for 2h in the buffer comprising 0.1 M sodium cacodylate (pH7.4), glutaraldehyde 2.5%, paraformaldehyde 2%, CaCl2 10 mM. To contrast the membrane structures, especially the ER, samples were stained with 2% osmium tetroxide and 0.8% potassium ferricyanide at 4°C overnight. After staining, samples were embedded in Spurr resin (EMS) in casting molds and polymerized at 70°C for 16 hrs. Fixed roots in spurr blocks were cut longitudinally into 90 nm thick sections with an EM UC7 ultramicrotome (Leica) and placed onto 200 mesh copper grids. Observations were carried out on a FEI TECNAI Spirit 120 kV electron microscope. To acquire sufficient data for plasmodesmata quantification, we captured TEM images on 5 to 15 cell walls in the meristem region per cell layer per root. Plasmodesmata were identified and counted manually while ImageJ software was used to measure the cell wall length. Plasmodesmata density was calculated by dividing the plasmodesmata numbers by the observation area (cell wall length x section thickness (90 nm)).

#### Serial block face scanning electron microscopy (SBF-SEM) sample preparation and imaging

4-day-old Arabidopsis seedlings were fixed for 2h at RT in 0.1M sodium cacodylate (pH 7.4) with 2.5 % glutaraldehyde, 2 % paraformaldehyde, and 10 mM CaCl2. First staining was performed overnight at 4°C in 0.1M sodium cacodylate (pH 7.4) with 2 % Osmium tetroxide, 3 % potassium ferricyanide, and 10 mM CaCl2. For the four additional staining, samples were subsequently stained for 1h in 1 % tannic acid in water, 1h in 2 % osmium tetroxide in water, 3h in uranyl acetate saturated in water, and finally 3h in 30mM aspartic acid (pH5.5). All the staining steps were performed at RT. Then, dehydration was done in gradient ethanol-water at 4°C and finally exchanged with ultrapure acetone. The substitution was done using a gradient of EPON812 in acetone. The seedlings were processed entirely, roots were cut out from the shoot at the last step only. Samples were embedded in 812 Epoxy resin (Agar Scientific) between aclar sheets with a 200 mm thick spacer and polymerized at 70°C for 16 h. Samples were mounted on aluminum pins using conductive silver epoxy resin (EMS). The acquisition was performed with 3View2XP system (Gatan) in GeminiSEM 300 (Zeiss) under high vacuum, 1.8kV acceleration voltage and using normal mode and 20mm aperture. Depending on the sample, the focal charge compensator was set to 70 to 90 %. Section thickness was set to 70 nm. The pixel size is 5 nm and the pixel time is 2 ms. Alignment and contrast normalization were done on Microscopy Image Browser. Data analysis was done using 3Dmod (IMOD). Plasmodesmata density was calculated similarly to TEM, except that plasmodesmata were counted every 4 sections within a single SBF-SEM stack. The cell wall surface was extracted from segmentation by IMOD (taking into account the sampling every four sections). For Fig.1, plasmodesmata density from SBF-SEM was applied to the whole wall surface extracted from a confocal z-stack of roots labeled with propidium iodide. Cell walls were segmented with Imaris and cell wall area was extracted with ImageJ.

#### Scanning transmission electron microscopy (STEM) tomography

4 days old Arabidopsis seedlings were fixed for 2h at RT in 0.1M sodium cacodylate (pH7.4) with 2.5 % glutaraldehyde, 2 % paraformaldehyde, and 10 mM CaCl2. Staining was performed overnight at 4°C in 0.1M sodium cacodylate (pH 7.4) with 2 % Osmium tetroxide, 1.5% potassium ferricyanide, and 10 mM CaCl2. The seedlings were processed entirely, roots were cut out from the shoot at the last step. Samples were embedded in 812 Epoxy resin (Agar Scientific) in casting molds and polymerized at 70°c for 16 hrs. 300 nm sections obtained with an EM UC7 ultramicrotome (Leica) are mounted on 100Cu grids coated with parlodion film and carbonated. 5 nm gold fiducials were placed on both sides of the sample. Acquisitions were done on ThermoFisher Talos F200S G2 STEM Unit and STEM-HAADF detector using the Fischione model 2045 tomography sample holder. Tilt series were reconstructed with Etomo (IMOD) software, using Hamming Filter at 50 and binning by 2.

#### Blue native gel electrophoresis and western blotting

2-gram 9-day-old Arabidopsis seedlings were ground in cold 8 ml vesicle isolation buffer (VIB) (0.45 M sucrose, 5 mM MgCl2, 1 mM dithiothreitol, 0.5 % polyvinylpyrrolidone, 50 mM HEPES, 1x Sigma protease inhibitor cocktail, 1x Roche complete Ultra protease inhibitor, 1mM PMSF, pH 7.5) for 30 minutes to fully break the cells and release the microsomes. Cell wall debris contains plasmodesmata and was separated from soluble contents by centrifuging at 1600xg for 20 min, 4 °C, and washed 3 times using VIB buffer. To solubilize the membrane protein in the cell wall, samples were resuspended in the 1 x NativePAGE sample buffer supplemented with 1 mM PMSF, 1x Sigma protease inhibitor cocktail (Sigma, P9599), non-ionic detergent 4% DDM (Life technologies, BN2005) or 2% digitonin (Merck, D141) and incubated for 1 h, at 4°C with gentle agitation. After solubilization, insoluble debris was removed by centrifugation (16000 x g, 20 mins, 4°C), and 20 μl of the supernatant was mixed with coomassie brilliant blue G250 (final concentration is 1/4 of the detergent in the sample) and loaded onto a 3-12 % NativePAGE Bis-Tris gel (Life Technologies, BN1001BOX). Gels were run under 55V for 2 h using dark blue cathode buffer (1x NativePAGE running buffer, 0.02% G-250) and then constant 2 mA for another 5 h using light blue cathode buffer (1x NativePAGE running buffer, 0.002% G-250). After running, the ladder was cut out and stained with Coomassie blue. The rest of the gel was wet transferred onto a PVDF membrane using NuPAGE transfer buffer (Life Technologies, NP0006-1) at 4°C for 20 h. After transfer, membranes were fixed in 8% acetic acid for 15 mins, washed twice with SDS buffer (62.5 mM Tris-HCl, pH6.8, 2% SDS) to expose the epitope for 30 mins, blocked with 5% non-fat milk in TBST and immunoblotted by monoclonal Anti-GFP (1:1500) (Sigma, Cat#Ref 11814460001, RRID: AB_390913) in the blocking solution.

#### SDS-PAGE and western blotting

Native extracted protein samples were mixed with 6x Laemmli buffer (Alfa Aesar, J61337, contains 9% SDS and 9% b-mercaptoethanol), denatured at 50°C for 30 mins, and then loaded onto TGX Stainfree 10% SDS-PAGE gels (Bio-rad, 1610185). Gels were wet-transferred to the PVDF membrane using CAPS buffer (2.21 g/L CAPS, 10% ethanol, pH 11) at 4°C overnight. After transfer, membranes were blocked with 5% non-fat milk in TBST and immunoblotted by monoclonal anti-GFP (1:1500) (Sigma, Cat#Ref 11814460001, RRID: AB_390913) in the blocking solution.

#### Co-Immunoprecipitation

The appropriate constructs were transiently expressed in *N. benthamiana* (see Transient expression in *N. benthamiana* section). Two days after agroinfiltration, approximately 0.5 g of tissue was collected per sample and immediately frozen in liquid nitrogen. Frozen tissue was grinded and mixed with 1 mL of protein extraction buffer (150 mM Tris-HCl, pH 7.5; 150 mM NaCl; 10 % glycerol; 10 mM EDTA, pH 8; 1mM NaF; 1 mM Na2MoO4; 10 mM DTT; 0.5 mM PMSF; 1% (v/v) P9599 protease inhibitor cocktail (Sigma); 1 % (v/v) Igepal), followed by incubation for 40 min at 4 C with continuous but gentle rotation. Protein extracts were centrifuge at 4C and 9000 g during 20 min and the supernatants were collected and filtered through Poly-Prep Chromatography Columns (#731-1550 Bio-Rad). 50 mL of each of the clear supernatants were kept as “input” samples and the rest were diluted with a washing buffer (same as extraction buffer but without Igepal) in a 1:1 ratio. Samples were then mixed with 15 mL of equilibrated GFP-Trap agarose beads and incubated at 4°C during 2 h with continuous but gentle rotation. Beads were then precipitated by centrifugation for 30 s at 500 g and washed 3 times with a washing buffer, followed by a final centrifugation for 30 s at 2000 g. Supernatants were discarded and the beads were mixed with 50 mL of Laemmli buffer and incubated for 20 min at 70°C. Finally, the beads were precipitated by centrifugation during 2 min at 2500 g and the supernatant with the eluted proteins were recovered. “Input” and “immunoprecipitated” samples were analyzed by western blot. For immunodetection we used monoclonal anti-mRFP (1:1000) (ChromoTek, clone 6G6, RRID: AB_2631395) and monoclonal anti-GFP (1:1500) (Sigma, Cat#Ref 11814460001, RRID: AB_390913).

#### Fluorescence recovery after photobleaching (FRAP)

FRAP experiments were performed on a Zeiss LSM 880 confocal microscope equipped with a Zeiss CPL APO x 40 oil-immersion objective (numerical aperture 1.3). YFP or GFP was excited at 488 nm with 1% argon laser power, and fluorescence was collected with the airyscan detector using BP495-550 + LP 525 filter. Photobleaching was performed on rectangular regions of interest (ROIs) at the cell plate of root epidermal meristem cells with the 488 nm excitation laser set to 100%. The FRAP procedure was the following: 5 pre-bleach images, 10 iterations of bleaching with a pixel dwell time set at 1.51l s, and then 50 images post-bleach with the “safe bleach mode for GaAsP”, bringing up the scan time up to approximately 2 s. The recovery profiles were background subtracted and then double normalized in the FRAP analysis website: https://easyfrap.vmnet.upatras.gr/?AspxAutoDetectCookieSupport=1

#### Fluorescence loss in photobleaching (FLIP)

FLIP experiments were performed on a Zeiss LSM 880 confocal microscope.5-7 days old HDEL-YFP or YFP-MCTP4 seedlings were stained with FM4-64 and root epidermal dividing cells were manually identified based on cell plate morphology. For the photo-bleaching and live imaging experiment, seedlings were placed above a ½ MS, 2% agarose solid medium to avoid stresses and drifting. A small ROI (1/5-1/10 of the bleaching daughter cell) was photobleached every 5 s using a 488 nm excitation laser, 100% power, and 20 iterations until the signal was no longer visible in the bleached daughter cells. 5 pre-bleaching and 55 post-bleaching images were taken using an Apochromat 40x/1.3 Oil DIC UV-IR M27 objective with a 488 nm laser and a 505-550 nm emission filter. Due to the growth of the roots, cells drifted in x-y and images were drift-corrected using ‘correct 3D drift’ in ImageJ. To analyze the signal loss over time, two daughter cells (one of which was selected for photo-bleaching), 3 surrounding reference cells, and 1 background position were manually outlined and their fluorescent signals were quantified at each time point and calculated as (mean gray value (bleached or unbleached daughter cell) - mean gray value (background)) / (mean gray value (3 ref cells) -mean gray value (background)). The average of the first 5 pre-bleaching images was set to 100 %, and all the following images during repeated bleaching were normalized accordingly. The measurements of all cells, at each stage of the cytokinesis, were combined and the average mean and SD were calculated and plotted using Prism9.

#### Correlative light and electron microscopy (CLEM)

For sample high-pressure freezing, cryo-substitution, microscope acquisition, and correlations we used the process described by (*37*). Briefly, root tips from 5-day-old *mctp3mctp4* complementation seedlings (*mctp3mctp4* x *pUBQ10-YFP-MCTP4*, line #4) were cut off and quickly high-pressure frozen in the 20 % BSA, followed by 30 h freeze-substitution (at -90°C) in acetone containing 0.1% uranyl acetate. After washing off the substitution mix, samples were embedded in HM20 Lowicryl resin. For acquiring the fluorescent signal from the sample, 150 nm sections were imaged with a 40x apochromatic N.A. 1.3 oil objective, 488 nm excitation laser, BP 495-550 filter, and airyscan detector from Zeiss LSM 880. Images were deconvolved using Zen Blue software to increase the signal/noise ratio and resolution. For acquiring TEM images, sections were observed by an FEI TECNAI Spirit 120 kV electron microscope. ec-CLEM software (https://icy.bioimageanalysis.org/plugin/ec-clem/) was used to analyze the correlation between TEM and light microscopy images based on the cell contours.

#### Electron tomography, tomogram reconstruction and image segmentation

For tomography done by an FEI TECNAI Spirit 120 kV electron microscope, we used the protocol described in (*26*). Sections were coated with 5 nm colloidal fiducials, which were used for image alignment. A series of tilt images were acquired over a -65 to 65 degrees range with an angular increment of 1 degree using FEI 3D explore tomography software. Tomograms were reconstructed using eTomo software (http://bio3d.colorado.edu/imod/). Segmentations were done by using 3dMOD (https://bio3d.colorado.edu/imod/doc/3dmodguide.html).

#### Confocal microscopy

All the confocal live-imaging experiments were recorded on a Zeiss LSM 880 equipped with 40x, 63x oil-immersion or 40x water immersion objectives and operated by ZEN Black 2011 software. Samples were labelled with 2mM FM4-64 for cell plate and plasma membrane staining and mounted between slide and coverslip with a thin layer of 1/2MS medium or on a Petri dish containing 1/2MS medium when imaged with a water objective. For detection of GFP or YFP fluorescence under the confocal mode, a 488 nm excitation laser and, 505-550 nm emission filter was used; for detection of tagRFP or mCherry fluorescence, a 556 nm excitation laser and, 570-625 nm emission filter was used; and for detection of FM4-64 fluorescence, a 556 nm excitation laser and 590-650 nm emission filter were used.

For airyscan imaging, detection of GFP or YFP was performed under airyscan Super Resolution mode, a 488 nm excitation laser and an emission band pass 495-550 were used. For the detection of FM 4-64, tagRFP or mCherry, a 561 nm excitation laser and emission 570-620 nm + long pass 645 nm were used. Images were taken sequentially frame when two colors were imaged at the same time to avoid signal cross-talk. Raw 2D and z-stack 3D images acquired with the airyscan detector were processed using Zen 3.3 blue.

#### Modified pseudo-Schiff propidium iodide (mPS-PI)

5-day-old Col-0, *mctp3mctp4*, *mctp3mctp4mctp6* seedlings were fixed and processed as in (*38*). Detection of propidium iodide was performed using a 514 nm excitation laser and 520-720 nm emission band pass. The entire root meristem was imaged with a z-stack of 1µm steps using LSM 880 confocal microscope.

#### Yeast

rtn1rtn2yop1spo7Δ yeast strain (*39*) was transformed with Pex30, AtMCTP3-TMR, AtMCTP4-TMR under the control of RTN1 promoter. Yeast strains were grown to mid logarithmic growth phase, serially diluted, spotted on to synthetic complete media plates, containing 2% glucose, 0.67% yeast nitrogen base without amino acids and an amino acids mix, with or without 5-FOA, and incubated at 30°C for 3 days. rtn1rtn2yop1spo7Δ strain with integrated ER marker ss-RFP-HDEL was transformed with AtMCTP3-TMR or AtMCTP4-TMR under the control of RTN1 promoter.

#### Sequence analysis

The prediction and delimitation of functional subdomains inside MCTP4 TMR was done by Hydrophobic Cluster Analysis (HCA) (*40*) and PSIPRED (*41*). The physicochemical properties and prediction of hydrophobic segments on alpha helices was done with HELIQUEST server (https://heliquest.ipmc.cnrs.fr/).

### Supplementary Text: Physical modeling of plasmodesmata formation

#### Essence of the physical model of plasmodesmata formation

Here, we propose a physical model of plasmodesmata formation. This is a theoretical model based on physical principles, which describes the morphology and energetics of plasmodesmata formation and stabilization, and the role that ER tubules and MCTP proteins play in this process.

The physical model serves to address two primary inquiries. Firstly, our ultrastructural analysis (Fig. 3) distinctly identifies two fenestrae populations differing in shape: those containing ER, with a minimum diameter of approximately 20 nm, and those lacking ER, which exhibit further constriction. We aim to investigate how the presence of ER tubules within constricting fenestrae may influence their final stabilization state. And secondly, we know from experimental data that MCTPs are important for the formation of plasmodesmata, induce high ER curvature and display membrane tethering activity. We aim at understanding the mechanical functions of MCTPs in influencing fenestrae to achieve the final stabilized state.

To test those questions on physical grounds, we developed a semi-quantitative equilibrium mechanical model to analyze physical mechanisms by which ER tubules and MCTP proteins could promote incomplete cytokinesis and bridge formation. In essence, the results of our model indicate that the presence of an ER tubule within a constricting cell plate fenestrae poses an extra challenge for fenestrae closure, as the tube needs to be either removed or severed before fenestrae closure (in physical terms, the presence of an ER tubule adds an extra energy barrier to the process of fenestrae closure, see discussion below). In addition, the presence of MCTP proteins with the capacity to tether the ER and the fenestrae membrane (referred after as the PM) and induce/stabilize ER membrane curvature contributes to the stabilization of the ER-cell plate membrane bridges and to hinder fenestrae closure (in physical terms, MCTPs add an additional energy barrier to fenestrae closure, see discussion below).

These types of physical models approach cell membranes as continuous surfaces that can be described by physical principles (such as lipid monolayer elastic theory). This modeling strategy yields valuable insights for our purposes: *(i)* it elucidates under which conditions incomplete cytokinesis and plasmodesmata bridge formation are energetically favored over complete cytokinesis (full fenestrae closure), thereby indicating the system’s inclination towards either outcome; *(ii)* it tracks morphological transitions from an initial state with widely open fenestrae and an ER tube to eventual ER tube removal and fenestrae closure; *(iii)* it quantifies the energy cost associated with the intermediate structures formed during these morphological transitions; and *(iv)* it identifies and quantifies the energy barriers that exist in the course of these transitions, barriers that can kinetically inhibit the cytokinesis process and, hence, promote the stabilization of plasmodesmata bridges (viewed as open cell wall fenestrae).

In developing such a physical model for plasmodesmata formation, two fundamental aspects must be considered: *(i)* a comprehensive description of the geometry of the system and potential shape transitions; and *(ii)* an understanding of the underlying physics governing the energies involved. The state of the system, whether it is in equilibrium or in a kinetically-arrested locally stable (i.e. metastable) state, is characterized by the concept of free energy, a fundamental thermodynamic quantity. Free energy encapsulates both the energetic contributions within the system and the entropic effects arising from their configurations. It serves as a state function to evaluate the stability of different structural configurations and to predict the transitions between them. By analyzing the free energy landscape with respect to relevant parameters, such as fenestrae diameter or the local concentration of MCTP proteins, we can discern the energetically favorable configurations (local minima of the free energy) and identify the barriers that hinder or facilitate transitions between them (this is what we computed for Fig. 3D, Fig.S5 and Fig. S15). Thus, a comprehensive understanding of both the geometric and energetic aspects of the system is essential to elucidate the mechanisms underlying plasmodesmata formation and stabilization. In the following paragraphs, we explain in lay terms both the geometry of the system and the underlying physics involved, before providing a more precise mathematical formulation in the following section.

#### Geometry of the system

The system is a single bridge (plasmodesma) of fixed length, *L*, that consists of a constricting cell wall fenestra and an ER tubule, which can contain a certain concentration of MCTP tethers, *ϕ*_*t*_. The ER tubule is modelled as a cylinder of radius, *R*_*t*_, connected to the ER tubular network (bulk ER) in both daughter cells. Based on ultrastructural analyses (this work and (*18*)), we restrict the geometry of the cell wall fenestrae to that of a cylindrical opening of length, *L*, and pore radius, *R*_*p*_. The length of fenestrae is kept constant in the model because experimentally we did not observe obvious differences between fenestrated sheet (FS) and cross wall (CWR) stage. A schematic representation of the system geometry is shown in Fig. S15A.

#### Underlying physics of the model

The free energy of the system arises from various factors, which we can categorize into those associated with (*1*) the fenestrae and surrounding cell plate, (*2*) the ER tubule, and (*3*) PM-ER interaction terms. Additionally, we consider (*4*) the effects of MCTP enrichment in the ER tube. We will now explain the essence of these contributions in lay terms, and in the following section we will derive the mathematical formulation of these contributions.

- First, the fenestra free energy reflects the thermodynamic work of creating a PM fenestra from a membrane reservoir. This thermodynamic work includes contributions from *(i)* the constricting force of cell plate expansion, promoting fenestrae closure; *(ii)* the turgor pressure, opposing fenestrae closure; and the elastic free energy of the PM, comprising *(iii)* resistance to membrane curvature (the so-called bending energy), which opposes fenestrae closure (especially for small fenestrae), and *(iv)* plasma membrane tension, promoting fenestrae closure (see Fig. 3E). The interplay between these factors determines the preferred equilibrium configuration of fenestrae in the absence of ER bridges (what we referred in the main text as the “no ER situation”).
- Second, the ER tubule energy represents the thermodynamic work associated with the remodeling (e.g., shrinking) of an ER tubule within a fenestra, still connected to the ER tubular networks of the two daughter cells, acting as membrane reservoirs. This energy includes the elastic energy of the ER membranes, with contributions from *(i)* bending energy of the ER tube (resisting tubule shrinking and hence impeding fenestrae closure), and *(ii)* ER membrane tension (promoting tubule narrowing and therefore not impeding fenestrae closure) (see Fig. 3E). Alone, ER tubule energy determines the tubule radius by the balance between bending and tension. However, when considering the overall system (ER tube, PM fenestrae, and possible interaction terms between them), the ER tubule energy influences the morphological fate of the system.
- Third, the interaction free energy is associated to interactions between the ER and the PM. This includes *(i)* hydration-mediated repulsion energy from the water layer between the membranes, preventing ER and PM fusion (see Fig. 3E); and *(ii)* MCTP-mediated tethering energy (adhesion energy) between ER and PM. For the latter, we assume a fraction of membrane tethers (the MCTP proteins) is present at the ER bridge. If the distance between the ER bridge and the enclosing fenestrae is small enough (≤10 nm based on (*18*)), MCTPs can tether the two membranes, stabilizing these contacts. In addition, we would like to mention that additional physical factors can control or modulate the spacing between closely apposed membranes. Besides the hydration-mediated repulsion considered in our model, other terms such as fluctuation-mediated steric repulsion interaction between membranes, Van der Waals attraction, or protein-protein interactions, might play a role. In here, for the sake of simplicity, we only considered a single term, the hydration-mediated repulsion, that prevents the collapse of the two membranes in apposition.
- And fourth, we consider physical factors associated with (or driving) potential enrichment of MCTP tethers in the ER bridge. Generally, enriching membrane proteins in local membrane subdomains, such as the ER bridge here, is entropically unfavorable, as free diffusion leads to a homogenous equilibrium distribution. However, factors such curvature-dependent sorting (when proteins generate or sense membrane curvature) and tethering-mediated sorting (when specific membrane subregions are primed for membrane-membrane interaction mediated by these proteins) can counterbalance those effects, driving local protein enrichment. These interactions create an imbalance in component diffusion into and out of the bridge, ultimately leading to protein enrichment. These factors need to be incorporated into our theoretical physical model by adding *(i)* an entropic free energy term penalizing component unmixing across the ER membrane; *(ii)* the tethering energy mentioned earlier as part of the interaction free energy; and *(iii)* a modification of the ER tube bending energy to account for the influence of MCTP concentration on the preferred (spontaneous) curvature of the ER tube.

In physics, energies are meaningful only when measured relative to a reference state, enabling the interpretation of the physical changes in terms of energy differences rather than of absolute values. Hence, we relate the total free energy of the system, *F*, to the free energy of a reference state, *F*_0_. In our case, we choose as the reference state a system formed by a fully closed fenestra and no ER tubule (akin to the complete cytokinesis state). In this case, the ER tubule is assumed reabsorbed to either of the two daughter cells, with a homogeneous bulk concentration of MCTP proteins, *ϕ*_*b*_, and acquire the preferred (mechanically stable) tube radius, 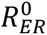. Our aim is to provide a mathematical formulation of the aforementioned physical mechanisms at play, enabling us to compute the free energy difference, Δ*F* = *F* − *F*_0_, as a function of the system and MCTP distribution along the ER membrane. This mathematical formulation of our physical model is outlined in the following sections.

Fenestra free energy

The fenestrae free energy is the thermodynamic work of creating a PM fenestra, and as outlined above, includes the following terms:

- *Constricting force of cell plate expansion.* We model the centrifugal expansion of the cell plate by implementing a general constricting force that acts towards fenestrae closure. We assume that such constricting force per unit length, *f*_*constr*_, is isotropically distributed all over the cell plate membrane and acts in the radial direction. Hence, the thermodynamic work done by such a force, ^Δ*F*^*constr*, reads as

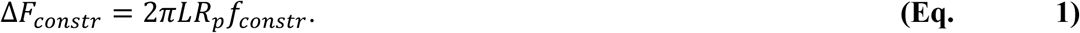
- *Turgor pressure.* Plant cells are under a high turgor pressure, Δ*p*, which contributes to the free energy as

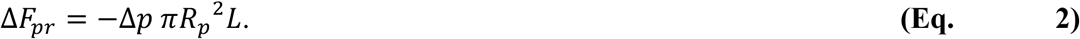
- *Bending energy of the PM pore.* The closure of the pore needs to overcome the bending energy of the pore rim. The free energy change is given by the Helfrich model of membrane curvature (*42*). Infinitesimally, the model reads as

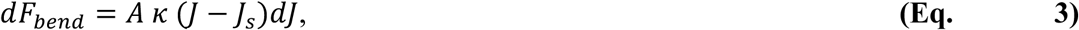 where *A* is the surface area of the membrane into consideration, ***κ*** is the bending rigidity, *J* is the total curvature (that is, the sum of the two principal curvatures), and *J*_*s*_ is the spontaneous curvature, which we consider to be zero at the PM. Hence, the total free energy change of creating a cylindrical pore of area 2*πLR*_*p*_, and radius *R*_*p*_ (and therefore total curvature *J* = 1/*R*_*p*_) out of flat membranes is 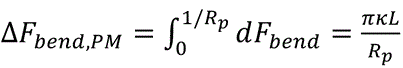. We will neglect the term coming from the Gaussian curvature, which reads as Δ*F*_*Gauss*_ = −4*π**κ***_*G*_, where ***κ***_*G*_ is the modulus of Gaussian curvature, as it is much smaller than the other energy terms. In addition, although experimentally measured values of the pore radius, *R*_*p*_, are relatively larger than the typical thickness of a lipid monolayer, *δ* ≈ 2*nm*, we not only take into account the curvature at the bilayer midplane, *J* = 1/*R*_*p*_, but consider each of the two monolayers independently (see ***Eq. 10*** below), yielding

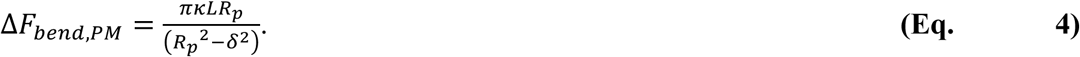
- *Plasma membrane tension energy.* The PM is under a certain lateral tension, *σ*_*PM*_, and hence the tension-associated free energy emanates from the change in the free energy of the reservoir. The change in the apparent area corresponds to the transition from a sealed fenestra of area 2*πR*_*p*_^2^(corresponding to the two circular membrane disks on each daughter cell) to a cylindrical fenestra of area 2*πR*_*p*_*L*, and hence the energy is

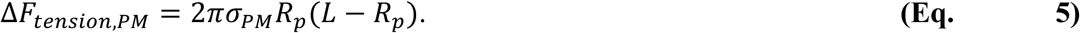

Hence, the total free energy per fenestrae is given by

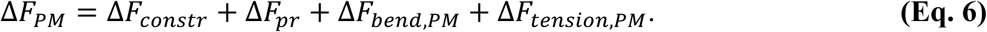

#### ER tubule energy

Next, we account for the changes in the free energy associated with the presence of a narrow ER tubule inside a fenestra. As before, we compute the free energy change required to generate a cylindrical ER tubule of radius *R*_*t*_, and length *L* out of the ER tubular network reservoir, where stable ER tubules have an equilibrium radius, *R*_*ER*_^0^ (see Fig. S15A). As outlined above, this energy includes two terms: a term associated to bending deformation of the tubule, and a term associated with the generation of the ER tubule against the membrane tension of the ER network reservoir. Following the classical approach to membrane thermodynamics and energetics (*43*, *44*), we split the thermodynamic work of ER tubule generation within a fenestra in two steps: first, the acquisition of membrane area out of the ER tubular network to generate the ER tubule within a constricted fenestra; and second, the bending deformation of such tubule.

- *ER membrane tension energy.* The ER network is under a certain lateral tension, *σ*_*ER*_, and hence the free energy change associated to the generation an ER tubule of surface area *A*_*t*_ = 2*πR*_*t*_*L* out of the ER membrane reservoir tension energy is

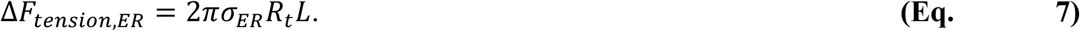

#### Bending energy of the ER tubule

Similar to the bending energy of PM fenestrae (***Eq. 3*-*4***), the bending energy change associated to the deformation of an ER tubule of area *A*_*t*_ = 2*πR*_*t*_*L* and an initial radius 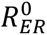 to a tubule of the same area and preferred curvature but with a different radius, 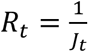, is given by

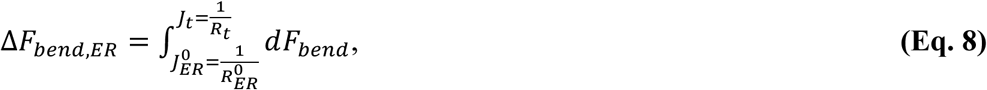

where, using the form for *dF*_*bend*_ given by (***Eq. 3***), and taking the bilayer midplane as the reference surface (of area *A*_*t*_), we get to

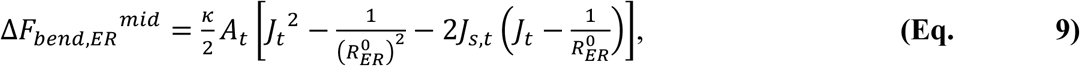

where the initial ER radius 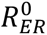 can be obtained as 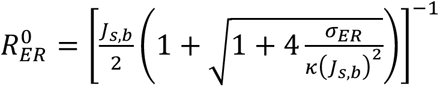 (*44, 45*); *σ*_*ER*_ is the lateral tension of the ER membrane; ***κ*** is the bending rigidity of the ER membrane (which we consider equal to that of the PM); *J*_*s*,*b*_ is the bare spontaneous curvature of the bulk ER tubular network; and *J*_*s*,*t*_ is the spontaneous curvature in the ER desmotubule, which can in principle depend on the concentration of curvature-inducing factors, such as MCTP proteins (see below). For very narrow membrane tubules, where the tubule radius, *R*_*t*_, is of comparable magnitude as to the monolayer thickness, *δ*, this equation has to be split in two parts, one for each of the monolayers (the bending energy of each monolayer is substantially different for radius of curvature of the order of the monolayer thickness (*46*). Using the equations for parallel surfaces (*47*), we can relate the total curvature and surface area of the two monolayer neutral surfaces (outer and inner) to that of the bilayer midplane (mid) as:

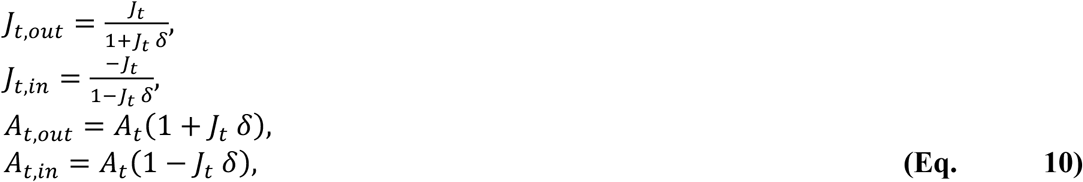

and, accordingly, split the bending energy in **(Eq. 9)** to the energies of each of the two individual monolayers as

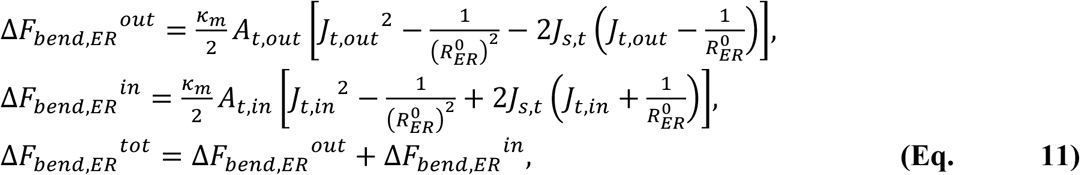

where ***κ***_*m*_ = ***κ***/2 is the monolayer bending rigidity, and we have taken into account the opposite sign of the curvatures in the inner monolayer.

Hence, the total free energy changed associated with the presence of a thin ER tubule within a PM fenestra is given by the sum of the terms in **Eqs. 7** and **11**,

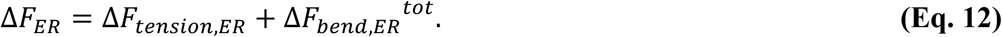

#### Interaction free energy

We next account for the energy terms resulting from the aforementioned interactions between the ER and the PM at fenestrae / nascent plasmodesmata:

- *Hydration energy.* Finally, we consider the hydration-mediated repulsion between polar lipid headgroups of opposed bilayers at the ER-PM interface (*48*, *49*). The free energy of hydration reads as

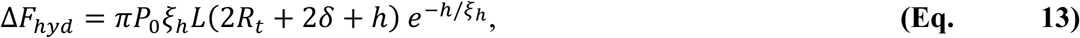 where *P*_0_ is the degree of membranes hydration, *ξ*_*h*_ is the characteristic hydration length, and *h* is the distance between the two membranes (the length of the water region). Because the *ξ*_*h*_ is of the order of a few Angstroms (see Table S1), the hydration energy is only relevant when the two membranes are in very close apposition to one another (*h* ≈ *ξ*_*h*_).
- *MCTP-mediated adhesion energy.* We consider that the MCTP tethers present at the ER tubule have an affinity to bind and anchor to the PM if the distance between the two membranes is small enough (in the range of the molecular size of MCTP proteins, i.e. (≤10 nm based on (*18*)). We can think of this tethering energy as an adhesion energy between the two membranes, which is then modelled as a simple step function,

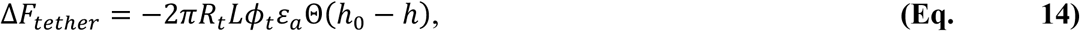 where *ɛ*_*a*_ is the binding free energy per MCTP and Θ(*x*) is the Heaviside step function, with *h* = *R*_*p*_ − *R*_*t*_ − 2*δ* being the distance between the two membranes, and *h*_0_the reach distance (the maximum tethering distance). The binding free energy, in case MCTP binding to the PM had a curvature dependency (curvature sensitivity) would read as *ɛ*_*a*_ = *ɛ*_*a*_^0^ − *⍺*_*J*_/*R*_*p*_, where *ɛ*_*a*_^0^ = *ɛ*_*a*_(*J* = 0), and *⍺*_*J*_ is the curvature sensitivity of the protein (*50*). For the sake of simplicity, we take *⍺*_*J*_ = 0 here.

Hence, the total interaction energy reads as

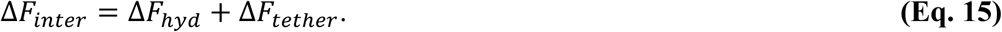

#### Free energy and effects of MCTP enrichment in the ER bridge

Next, we describe mathematically the physical factors associated with the potential enrichment of MCTP tethers in the ER bridge. This includes curvature-dependent and tethering-mediated sorting mechanisms, and necessitate the incorporation or modification of extra terms into our theoretical model.

#### Enrichment of MCTP in the cytoplasmic bridge

When molecular components, such as MCTPs, are non-homogeneously distributed across a continuous membrane, one needs to consider the free energy associated with such enrichment. This is basically an entropic free energy term, and, for the sake of simplicity, we ignore any enthalpic contribution of lateral self-association (oligomerization) of MCTP proteins. Following the derivation in (*44*), the entropic free energy associated with an enrichment of molecules (from a bulk concentration of *ϕ*_*b*_ to a local concentration in the tube, *ϕ*_*t*_) in a membrane patch of area *A*_*t*_ = 2*πR*_*t*_*L*, corresponding to the ER tubule area, reads as

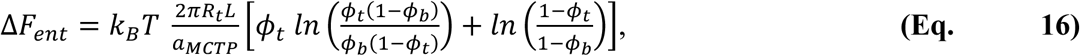

where *k*_*B*_*T* = 4.11 *pN nm* is the thermal energy, and *a*_*MCTP*_ is the cross-sectional surface area of a MCTP protein on the membrane plane. In addition, we could also include the term associated to the reduction in the entropy of the tethers due to confinement in the narrow space between the ER and PM, but we ignore this term for the sake of simplicity.

Finally, we describe how curvature generation by MCTPs alters the ER membrane energy. Enrichment or depletion of MCTPs would affect the mechanical equilibrium of ER tubules. To model this effect, we consider that MCTPs, as other membrane inducers, can be described by their effective molecular spontaneous curvature, *ζ*_*MCTP*_. Hence, we can express the spontaneous curvature of the bulk ER network as

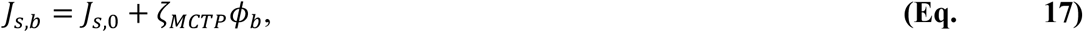

which has a dependence on the bulk ER network concentration of MCTP proteins, *ϕ*_*b*_, and where *J*_*s*,0_is the spontaneous curvature of the bulk ER tubular network in the absence of MCTP tethers. Similarly, the spontaneous curvature of the ER bridge is

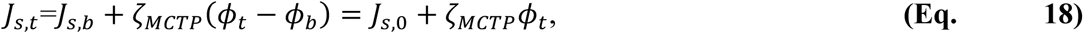

which also has a linear dependence on the area fraction of MCTP proteins on the ER tube, *ϕ*_*t*_, the proportionality factor, *ζ*_*MCTP*_, being the effective bilayer spontaneous curvature induced by MCTPs (see e.g., (*50*)). These two expressions are then introduced in the expression for the bending energy of the ER bridge, **Eq. 11**.

#### Total free energy of the system

When taking all these terms into account, the free energy of the system, including the cell plate fenestra (**Eq. 6)** and ER (**Eq. 12**) energies, the interaction terms (**Eq. 15**) and the entropic contribution of a possible MCTP enrichment (**Eq. 16**), the total free energy change of the system, Δ*F*, is

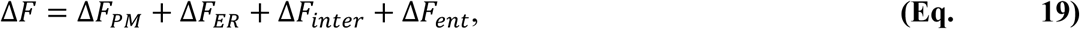

where the ER bending energy depends on the concentration of MCTPs, *ϕ*_*t*_, and therefore the total free energy change of the system is a function of tether concentration and the system geometry, represented by the fenestra radius, *R*_*p*_, and the ER tube radius, *R*_*t*_. Hence, we can write that Δ*F* = Δ*F*Y*R*_*p*_, *R*_*t*_, *ϕ*_*t*_Z. Besides, our model depends on a number of parameters, which have been experimentally measured or estimated (see Table S1 for details).

#### Analysis of the effects and relative contributions of individual free energy terms

The total free energy of the system comprises multiple terms (**Eq. 19**), some facilitating fenestrae shrinkage while others impede it. In this section, we illustrate how these different terms play contrasting roles in the process of plasmodesmata formation and discuss their respective contributions to the total free energy of the system. For the sake of simplicity of this illustration, we stick to the case with no MCTPs. The effects of including MCTPs in the system are discussed in the *Strategy of computations* section below. As the number of model’s parameters is large, we will use throughout this section the values stated in Table S1, unless otherwise stated.

First, the fenestrae free energy contains four contributions, as mentioned above (**Eq. 6**). Three of them – the turgor pressure, the PM tension, and the bending energy of the fenestrae PM – increase with decreasing fenestrae diameter, therefore impeding fenestrae shrinkage (Fig. S5A). Of these three, the PM tension plays a relatively minor contribution, while the bending energy dominates for small fenestrae sizes but remains negligible for large fenestrae sizes. The fourth term, the constricting force of cell plate expansion, facilitates fenestrae closure as it decreases with decreases fenestrae size. The combination of all these terms gives rise to a non-monotonous function that has a local minimum at a small fenestrae diameter, which corresponds to a locally metastable state (Fig. S5A).

Second, the free energy of the ER tubule has two contributions (**Eq. 12**): the tension of the ER membrane, promoting tube shrinkage, and the bending rigidity of the ER tube, which is a parabolic equation with a minimum at a certain tube size (Fig. S5B). Of note, this energy is solely the energy of the ER tubule, without taking into account any interaction terms with the membrane of the fenestrae (such as tethering or hydration energies).

The next step is to combine these two free energies and see how the effect of having an ER tube inside a closing fenestra hampers the abscission. When combining these energies, we also need to include the interaction term, in this case simply the hydration energy (**Eq. 13**). As now we have an extra free parameter, that is the distance between the membranes, *h*, we need to undergo a second optimization process (see section *Strategy of Computations* for details). Our model’s results indicates that, as the cell plate is maturing and fenestrae contract, the free energy of the system is totally dominated by the fenestrae energy term. As the size of the fenestrae reaches sizes of the order of the ER tubule radius, then the free energy of the tubule dominates the total free energy of the system and arrests the contraction creating a metastable state (Fig. S5C). Importantly, the interaction energy (the hydration energy) is negligible in the inter-membrane distance optimized case, as it prevents the two membranes from contacting one another, and sets the optimal intermembrane distance, *h*^∗^, to values between 1 and 2 nm (Fig. S5D). In summary, these plots allowed us to compute the optimal geometry of the system as the fenestrae is closing. In the initial steps of fenestrae shrinkage, the size of the ER tubule is not affected, as the intermembrane distance is large and there is no interaction between these two membranes. Further closure of the fenestrae beyond ∼32 nm in radius starts inducing a reduction of the ER tubule size due to the close contact (below 10 nm) between these two membranes (Fig. S5E).

#### Strategy of computations

Once we have the total free energy of the system (**Eq. 14**, or **Eq. 19** if including the effects of MCTP proteins) and the geometry of the system (Fig. S15A), we can study how the free energy changes as a function of the free parameters of the model (see Table S1). In particular, the model has two morphological free parameters, the fenestrae radius, *R*_*p*_, and the ER tubule radius, *R*_*t*_, and, if considering MCTP proteins, the concentration of MCTP on the ER tubule, *ϕ*_*t*_. In the case where we consider a naked fenestra (no ER tubule), there is only one free parameter, the fenestra radius, *R*_*p*_. Because the radii are measured at the bilayer midplane, the condition of non-overlapping membranes can be expressed as *R*_*p*_ > *R*_*t*_ + 2*δ*. Hence, because the distance between the two monolayers is *h* = *R*_*p*_ − *R*_*t*_ − 2*δ* > 0, we also use this parameter instead of *R*_*p*_, so, in the most general case (***Eq. 19***), we have Δ*F* = Δ*F*(*R*_*t*_, *h*, *ϕ*_*t*_). We then numerically find the set of free parameters {*R*_*t*_, *h*, *ϕ*_*t*_} that correspond to local minima of Δ*F*, and that are subject to the constraints given by *R*_*t*_ > *δ*;*h* > 0; 0 ≤ *ϕ*_*t*_ ≤ 1. Regarding the latter constraint, we will limit the maximum coverage of MCTPs on the ER tube to 50% (0 ≤ *ϕ*_*t*_ ≤ 0.5). On the one hand, because it is possibly not feasible to have larger protein area fractions due to the size of their cytosolic regions, and on the other hand because for large values of *ϕ*_*t*_, the entropic part of the free energy is not fully realistic (the entropic part of the free energy assumes relatively small protein enrichment). In addition, when alternatively considering the fenestra radius instead of the intramembrane distance, the non-overlap constraint is *R*_*p*_ > 2*δ*.

In the situation where we do not consider the effects of MCTP proteins, we proceed our optimization problem using a two-step approach. First, for any given value of the fenestra radius, *R*_*p*_, we start by finding the value of the ER tubule radius, *R*_*t*_^∗^Y*R*_*p*_Z, in the local vicinity of 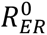, that minimizes the total free energy (***Eq. 14***). We then plot and minimize Δ*F*Y*R*_*t*_^∗^Y*R*_*p*_Z, *R*_*p*_Z with respect to *R*_*p*_, yielding Δ*F*_*opt*_ = Δ*F*Y*R*_*t*_^∗^Y*R*_*p*_^∗^Z, *R*_*p*_^∗^Z, where *R*_*p*_^∗^is the value of *R*_*p*_(in the region of small values of *R*_*p*_) that locally minimizes the total free energy (see Fig. 3D).

Similarly, when we include the effects of MCTP proteins into our model, we proceed our optimization problem using a two-step approach. First, for any given value of the fenestra radius, *R*_*p*_, we proceed with a two-parameter local minimization (using the “FindMinimum” or “NMinimize” functions in Wolfram Mathematica 9 software) to find the values of the ER tubule radius, *R*_*t*_^∗^Y*R*_*p*_Z, in the local vicinity of 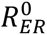, and of the MCTP area fraction, *ϕ*_*t*_^∗^Y*R*_*p*_Z, that together minimize the total free energy (***Eq. 19***). We next plot and minimize Δ*F* >*R*_*t*_^∗^Y*R*_*p*_Z, *R*_*p*_, *ϕ*_*t*_^∗^Y*R*_*p*_Z) with respect to *R*_*p*_, yielding Δ*F*_*opt*_ = Δ*F* >*R*_*t*_^∗^Y*R*_*p*_^∗^Z, *R*_*p*_^∗^, *ϕ*_*t*_^∗^Y*R*_*p*_Z), where *R*_*p*_^∗^ is the value of *R*_*p*_ (in the region of small values of *R*_*p*_) that, together with *R*_*t*_^∗^Y*R*_*p*_^∗^Z and *ϕ*_*t*_^∗^Y*R*_*p*_^∗^Z, locally minimizes the total free energy. For the results shown in Fig. S15B, we have optimized Δ*F* with respect to *R*_*p*_ and *R*_*t*_, and left *ϕ*_*t*_ as a free parameter to illustrate how the free energy of the system has the tendency to decrease with the enrichment of MCTPs in the ER tube.

#### Constriction force analysis

The constriction force provides a means to stabilize finite-size fenestrae. Indeed, in the absence of a constriction force, the turgor pressure would favor the opening of the fenestrae. From the fenestra energy, ***Eq. 6***, we can see that a locally stable solution at a finite pore size exists for constriction forces larger than a critical constriction force, *f*_*constr*_^∗^, which can be found as the solution to 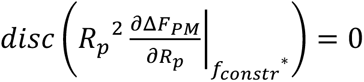, where *disc* is the polynomial discriminant. This gives the value of the critical constriction force as 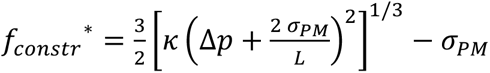. For the parameters in Table S1, we obtain *f*_*constr*_^∗^ = 2.1 *pN nm*. At this critical constriction force, the locally stable fenestrae radius, *R_p_*^∗^, is given by the solution to 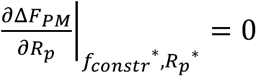, which is 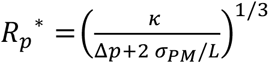. For typical parameters (see *Table S1*), one obtains *R*_*p*_* = 7.4 *nm*.

**Fig. S1.**
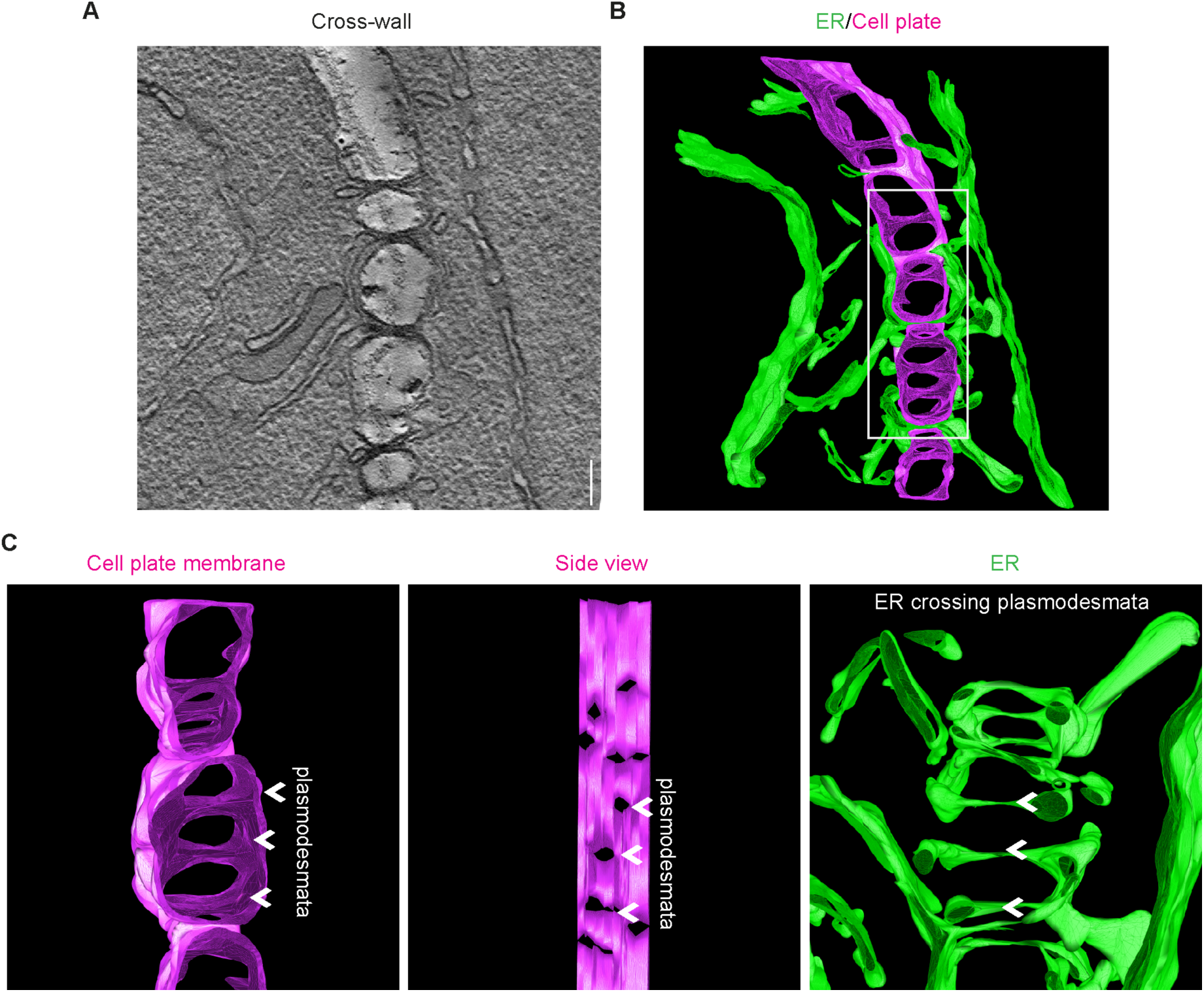
STEM tomography on cross-wall plasmodesmata bridges. **(A)** Reconstructed scanning transmission electron tomography of a cell plate segment at cross-wall stage in root endodermis meristem. **(B-C)** 3D segmentation of the tomography presented in (A). Top and side view of plasmodesmata embedded within the cell plate (C). ER is labeled in green and the cell plate membrane is labeled in magenta. Scale bar, 200 nm.

**Fig. S2.**
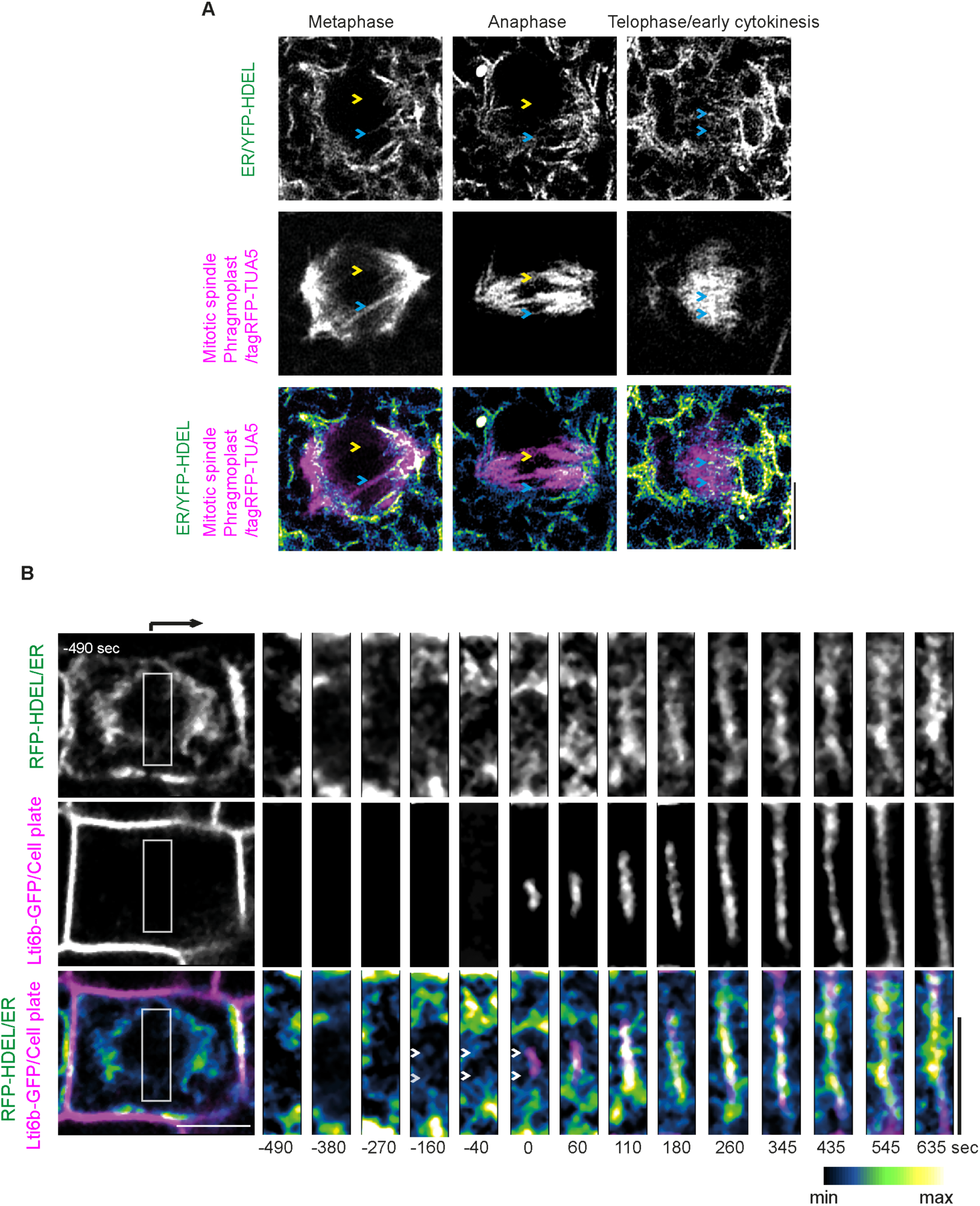
ER dynamics during mitosis and cytokinesis in epidermal cells of *A. thaliana* root. **(A)** Airyscan imaging of live root epidermal cells at three representative mitosis stages expressing ER (YFP-HDEL) and microtubule (tagRFP-TUA5) markers. Yellow arrows mark the ER staying away from the spindles at metaphase and anaphase. Blue arrows mark occasional signal overlapping between the ER and the spindles at metaphase and anaphase. The ER starts to accumulate at the future division plane at telophase/early cytokinesis. Representative of more than six cells from two technical replicates. **(B)** Dividing root epidermal cell expressing ER (RFP-HDEL) and plasma membrane (PM)/cell plate (Lti6b-GFP) markers, undergoing telophase (-490 s to 0 s) and cytokinesis (0 s to 635 s). Time 0 corresponds to the first appearance of the cell plate (Lti6b-GFP) signal. White arrowheads mark the ER that precedes the cell plate initiation. Representative of observations from 20 seedlings across six technical replicates. Scale bars, 5 μm.

**Fig. S3.**
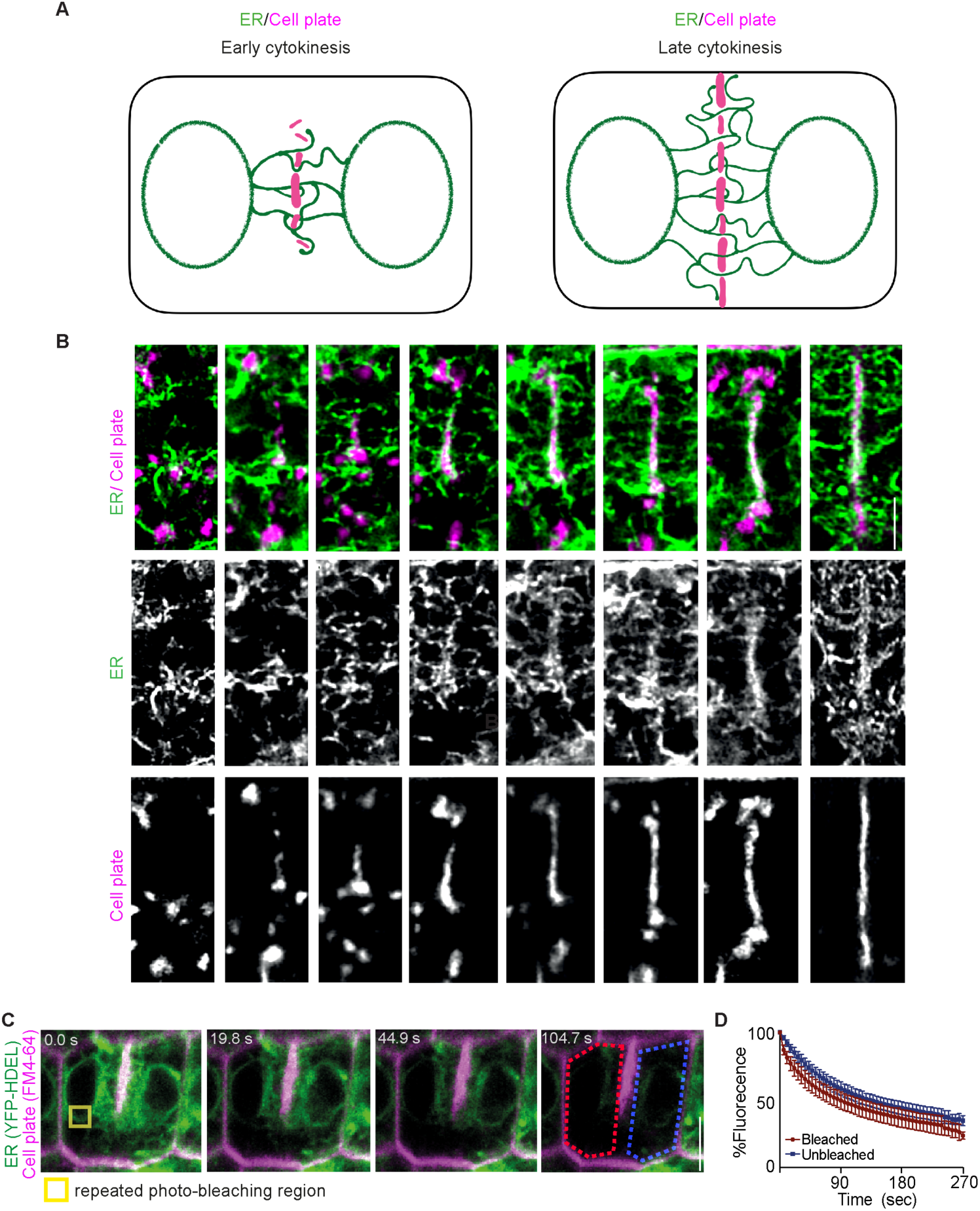
The ER is maintained as one continuous membrane network during cytokinesis spanning across the cell plate. Epidermal cells of *A. thaliana* root meristematic region. **(A)** Schematic illustration of the ER and cell plate interlinked association at the early and late cytokinesis as seen in (B)**. (B)** Airyscan imaging of root epidermal dividing cells showing representative ER-cell plate co-assembly starting from cytokinesis initiation until the end of the process. ER is labeled by YFP-HDEL (green) and the cell plate by FM 4-64 (magenta). **(C and D)** fluorescence loss in photobleaching (FLIP) on root epidermal cytokinesis cells expressing YFP-HDEL ER luminal marker and co-labeled by FM 4-64 to visualize the cell plate and the plasma membrane (PM). Note the simultaneous loss of the YFP-HDEL fluorescence in both daughter cells, while only one (outlined by red) is repeatedly photobleached. The bleaching region is outlined in yellow. Non bleached daughter cell is outlined in blue. Average fluorescence intensity at each time point, in bleached (red) and un-bleached (blue) daughter cells. The bars indicate the mean and SD. (n = 17 cells from 4 technical replicates). Scale bars, 5 μm (B); 2 μm (C).

**Fig. S4.**
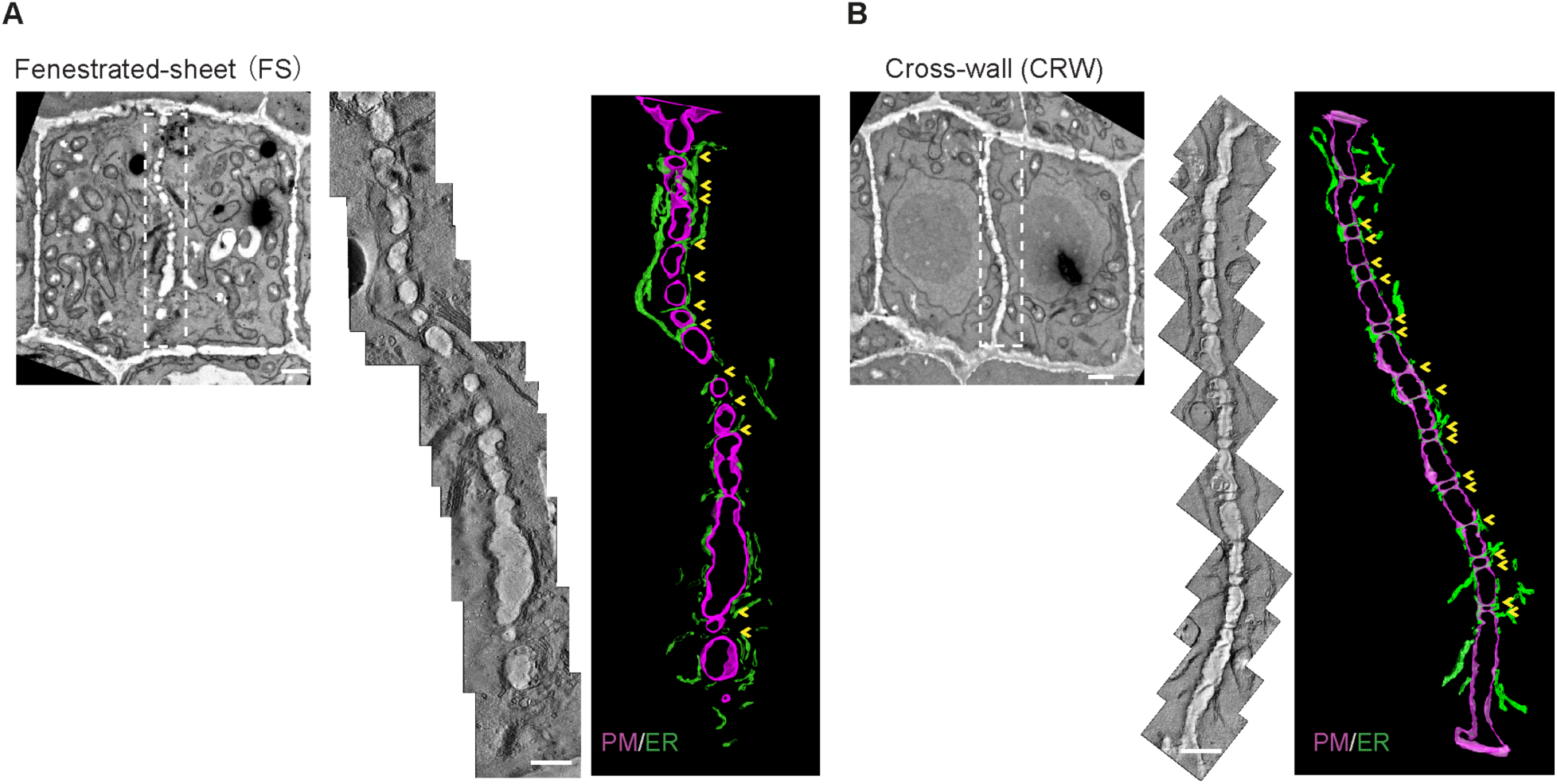
Electron tomography on cytokinesis cell plates at fenestrated-sheet (FS) and cross-wall (CRW) stages. Overview of the cell plate at fenestrated-sheet **(A)** and cross-wall **(B)** stages. Multiple electron tomography acquisitions were performed to cover the entire cell plate. Representative 2D images from each tomography were stitched together to illustrate the structural details of the cell plate organization and were manually segmented. ER is labeled in green and cell plate membrane (future plasma membrane PM) is labeled in magenta, fenestrae are indicated by yellow arrows. Scale bar, 1 μm (cell overview) and 500 nm (stitched cell plates).

**Fig. S5.**
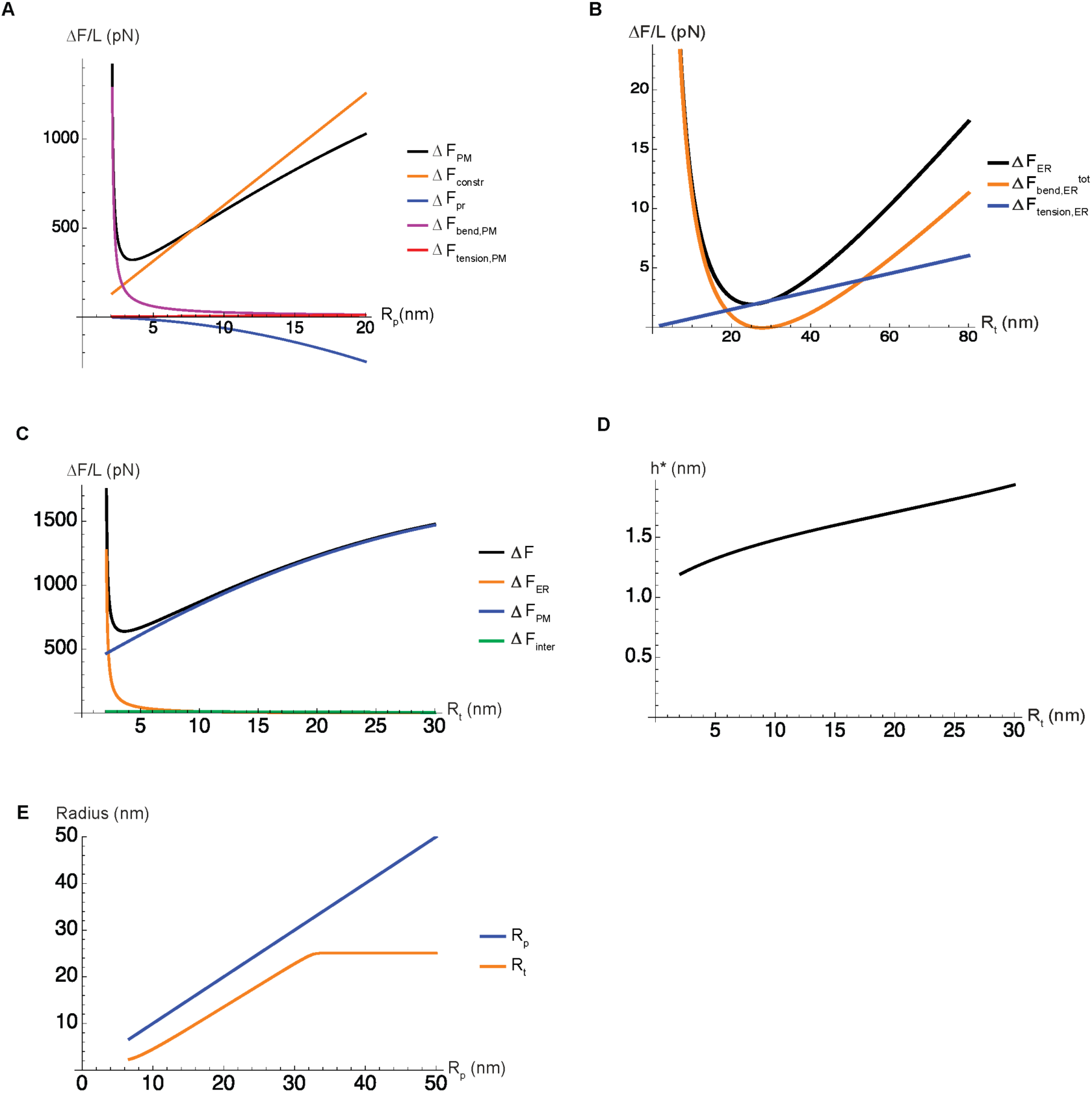
Analysis of the effects and relative contributions of individual free energy terms. The results of our free energy model (in the no MCTP situation) have been split and individually plotted for the different energetic contributions: **(A)** fenestrae free energy, **(B)** ER tubule energy, and **(C)** total free energy, including the interaction (hydration) energy. **(D)** Results of the optimization of the intermembrane distance (distance between the membranes of the ER and PM). **(E)** Correlation between the radius of the fenestra (*R*_*p*_) and of the ER tubule (*R*_*t*_) in the course of fenestrae closure (changes in the size of the fenestra radius).

**Fig. S6.**
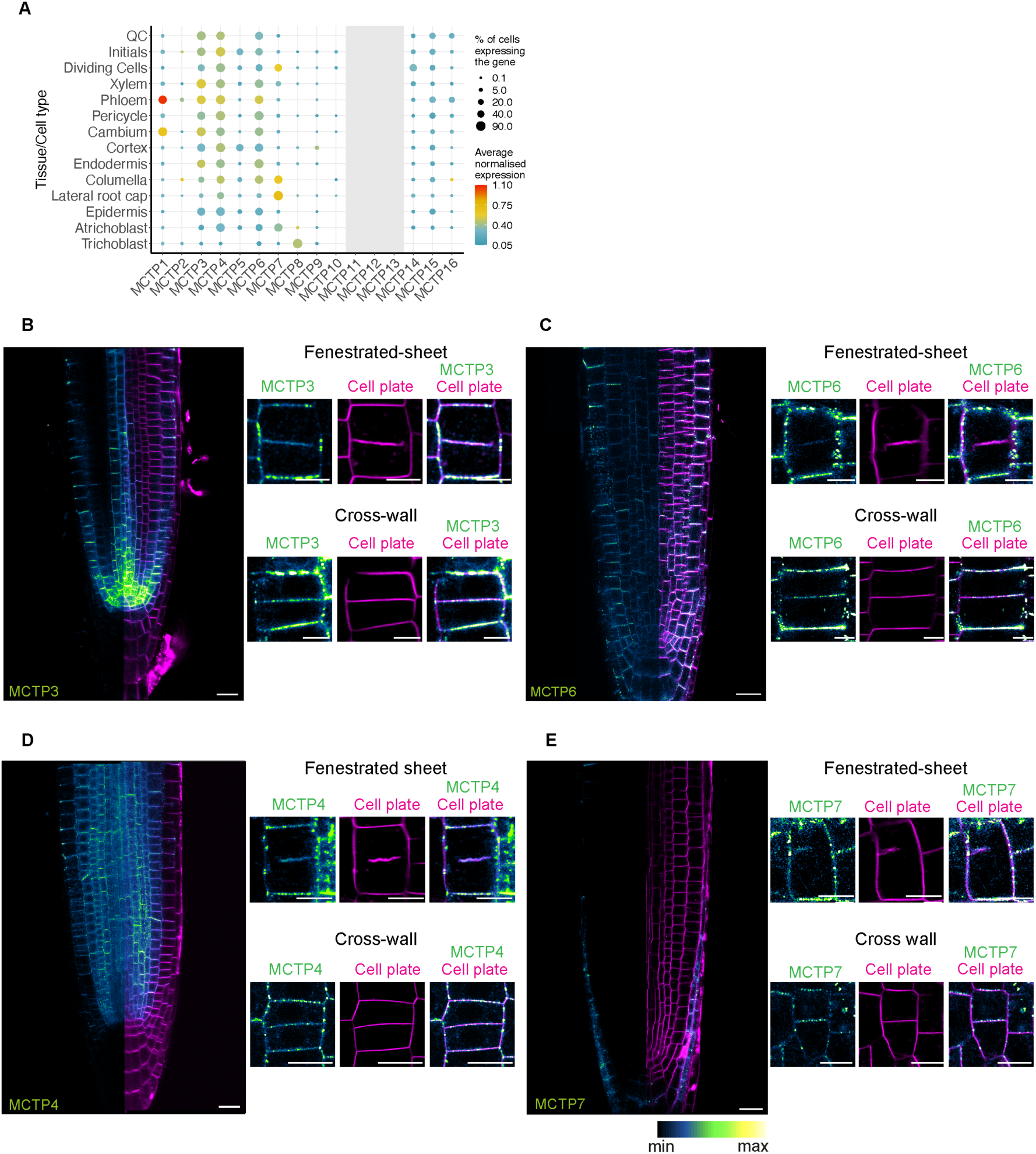
MCTPs expression pattern and subcellular localization during cytokinesis. **(A)** Dot plot showing the expression pattern of MCTP1-16 across different cell types in the *A. thaliana* primary root. The size of the circles represents the percentage of cells with expression, whereas the color indicates the scaled average expression. MCTP11, 12, 13 are not detected and marked as grey. (Based on single cell sequencing data obtained from (*26*). **(B-E)** Expression pattern of *pMCTP3-YFP-MCTP3* (in *mctp3* mutant background), *pMCTP4-YFP-MCTP4* (in *mctp4* mutant background), *pMCTP6-MCTP6-YFP* (Col-0 background) and *pMCTP7-YFP-MCTP7* (Col-0 background) in the roots with representative localization pattern in cells at fenestrated-sheet and cross-wall stage. MCTPs in green and plasma membrane (PM)/Cell plate (stained by FM4-64) in magenta. Expression and subcellular localization are consistent between n = 5 (MCTP3), n = 4 (MCTP4), n = 2 (MCTP6) and n = 4 (MCTP7) independent transgenic lines. Scale bars, 10 μm (root expression panels) and 5 μm (cytokinetic cell panels).

**Fig. S7.**
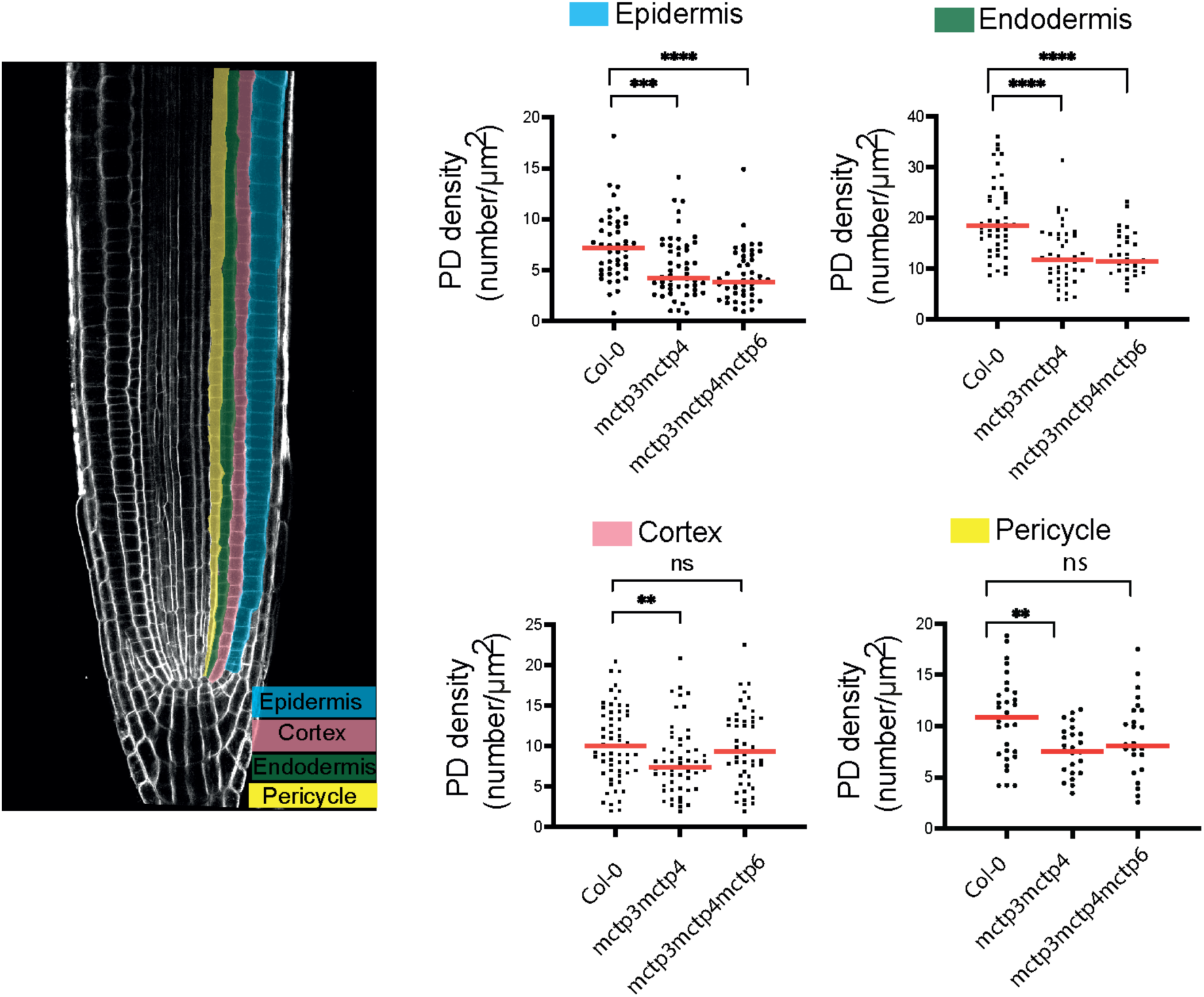
Plasmodesmata quantification in the *A. thaliana* wild-type (Col-0) and *mctp* mutants. Quantification of the plasmodesmata (PD) densities on post-cytokinetic wall (apico-basal walls, number of PD/μm^2^) in four cell types in the root meristem using transmission electron microscopy. The bars indicate the mean. Significance was tested using ordinary two-tailed Mann-Whitney U-tests (****, P<0.0001). Epidermis: n = 46 (Col-0), n = 53 (*mctp3mctp4*), n = 50 (*mctp3mctp4mctp6*) cells. Endodermis, n = 45 (Col-0), n = 45 (*mctp3mctp4*), n = 34 (*mctp3mctp4mctp6*) cells. Cortex n = 60 (Col-0), n = 55 (*mctp3mctp4),* n = 51 (*mctp3mctp4mctp6*) cells. Pericycle n = 31 (Col-0), n = 28 (*mctp3mctp4*), n = 27 (*mctp3mctp4mctp6*) cells from 5 roots of Col-0 and mutants.

**Fig. S8.**
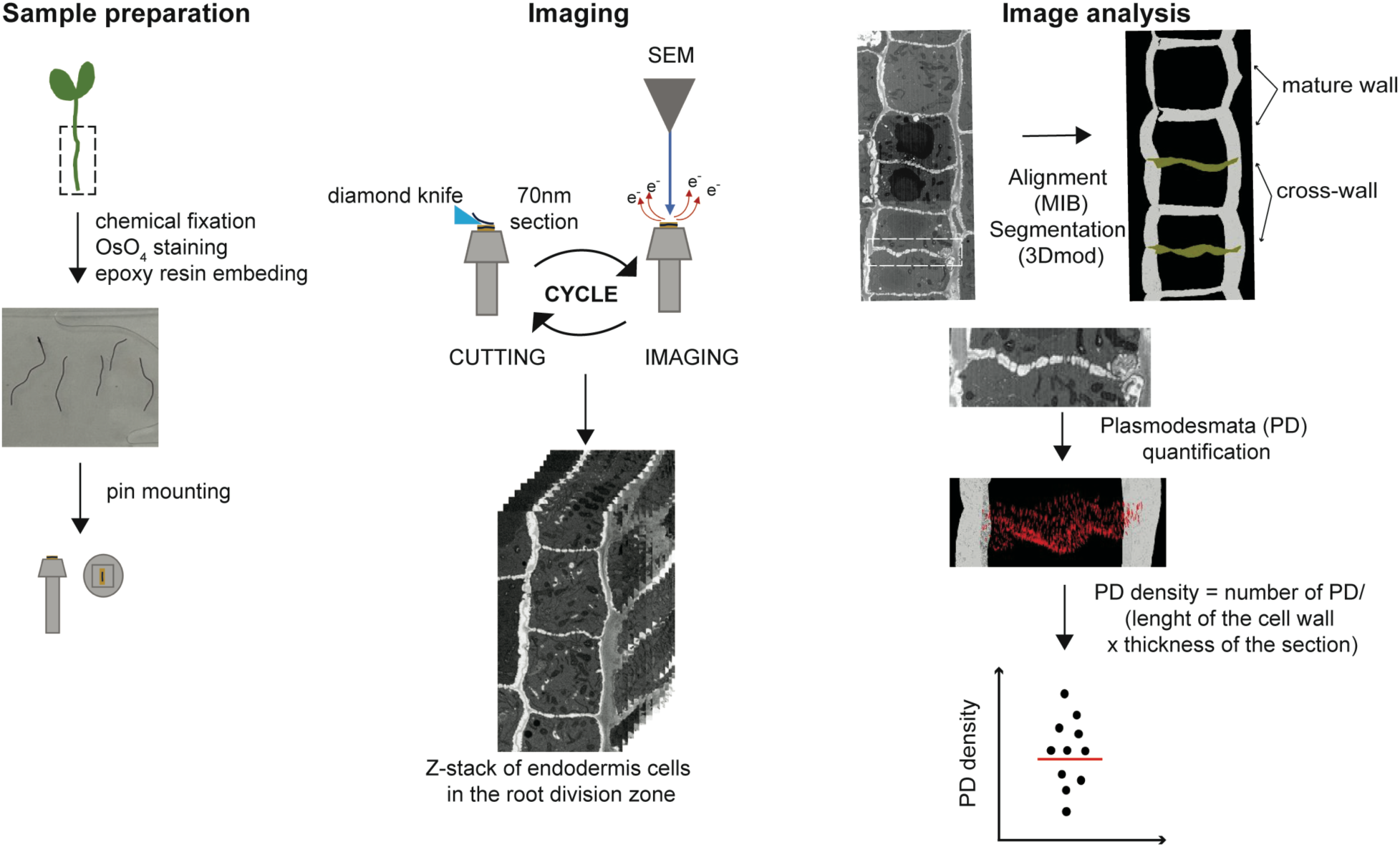
Workflow for plasmodesmata quantification from SBF-SEM imaging. SBF-SEM workflow is divided into three main stages: Sample preparation, imaging and image analysis. 4-day-old Arabidopsis seedlings are chemically fixed and highly stained with osmium tetroxide to enhance ER contrast. Root samples are embedded in epoxy resin, mounted on the top of aluminum pin, and trimmed to reach the region of interest: endodermal cells of the meristematic zone. The root is imaged in a high vacuum SEM chamber in a cycle as follows: the surface of the meristematic zone root is scanned and then a diamond knife cuts 70nm sections to obtain a z-stack from the sample surface to depth. The image z-stack is aligned (MIB software) and the ER, cell plate and plasmodesmata are annotated and segmented (3DMOD software) at cross-wall and mature wall stage for 3D visualization. From these data, plasmodesmata density was calculated as follows: number of PD/ (length of the cell wall x thickness of the section).

**Fig. S9.**
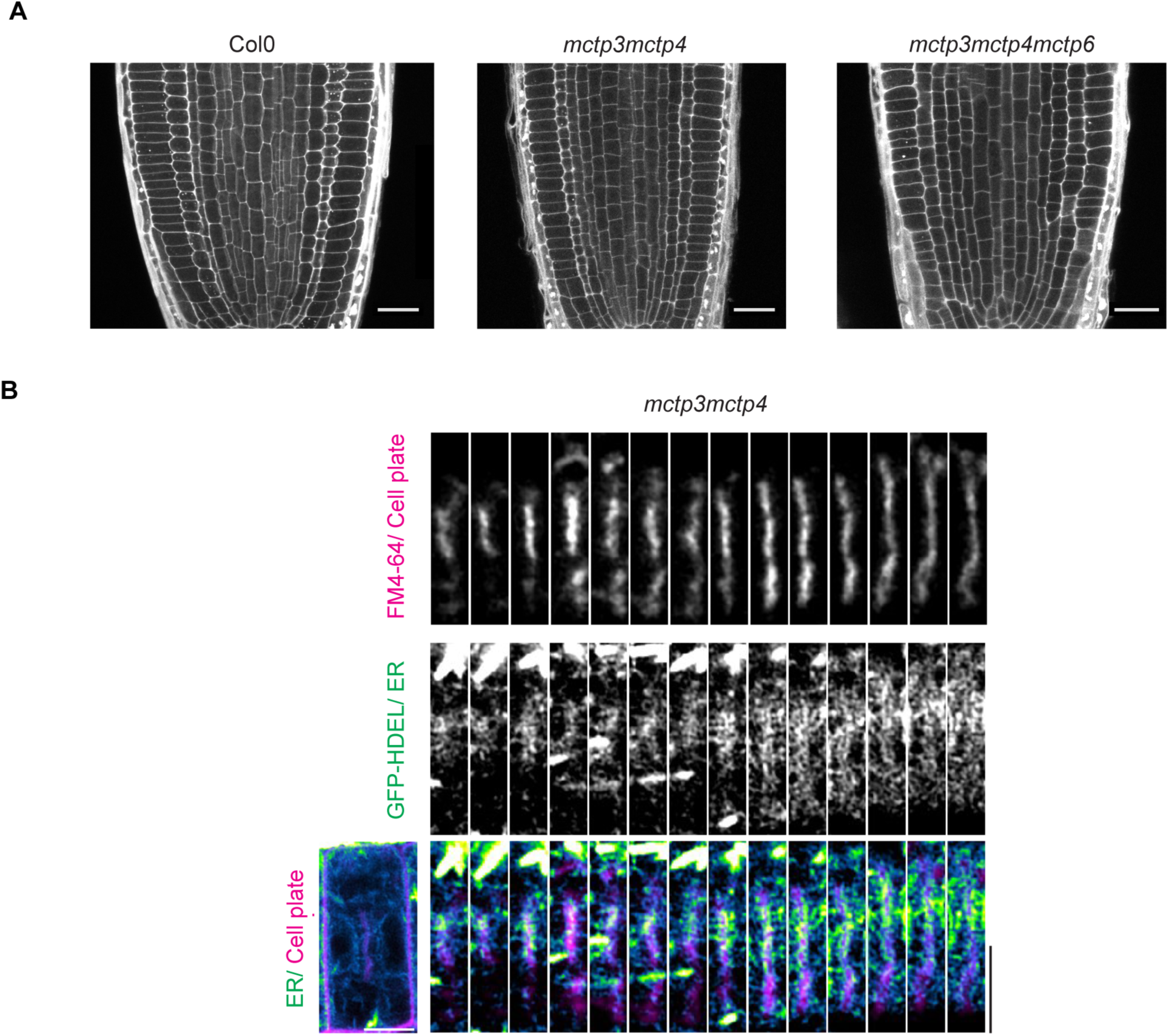
Cell wall staining and ER dynamics during cytokinesis in *mctp* mutants. (**A**) Pseudo-Schiff propidium iodide wall staining of Col-0, *mctp3mctp4* and *mctp3mctp4mctp6* root meristematic region. n = 26 for Col-0, n = 9 for *mctp3mctp4* and n = 11 for *mctp3mctp4mctp6* from 3 technical replicates. Scale bars, 20μm. (**B**) Time lapse of ER dynamics during cytokinesis by airyscan imaging of live root epidermal cells of *mctp3mctp4* mutant expressing GFP-HDEL. Representative of observations from 6 seedlings across 3 technical replicates. Scale bars, 5μm.

**Fig. S10.**
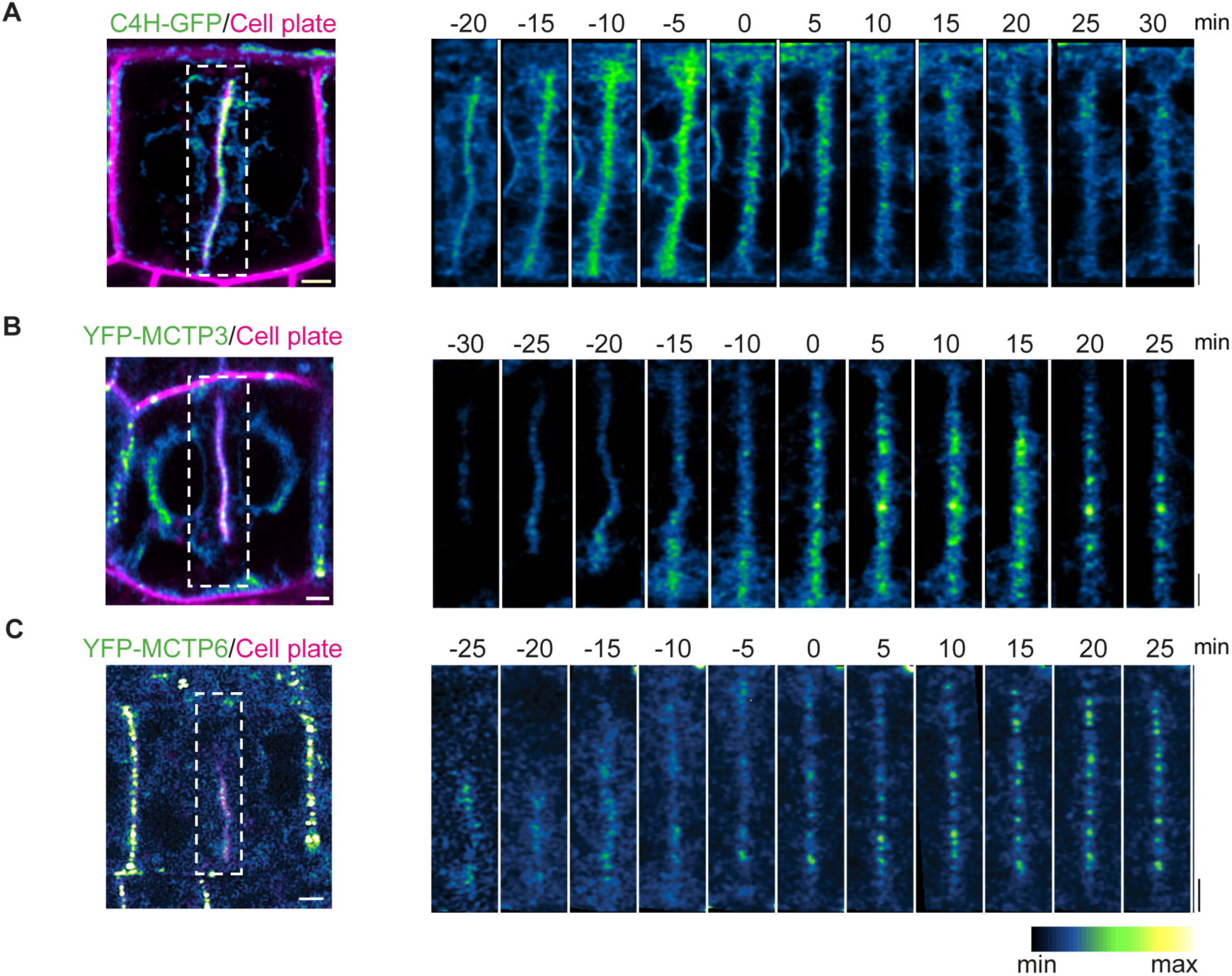
MCTPs accumulate at nascent plasmodesmata. **(A)** ER marker C4H-GFP (Col-0 background), **(B)** YFP-MCTP3 (*mctp3mctp4* complemented plants) and **(C)** YFP-MCTP6 (*mctp3mctp4mctp6)* complemented plants time lapse during cytokinesis, root epidermal cells. Time 0 indicates the start of the cross-wall stage. Note that MCTP3 and MCTP6 signals start to accumulate at the onsite of cross-wall stage. C4H-GFP, YFP-MCTP3, and YFP-MCTP6 in color coded green fire blue and cell plate in magenta. Scale bar, 2μm

**Fig. S11.**
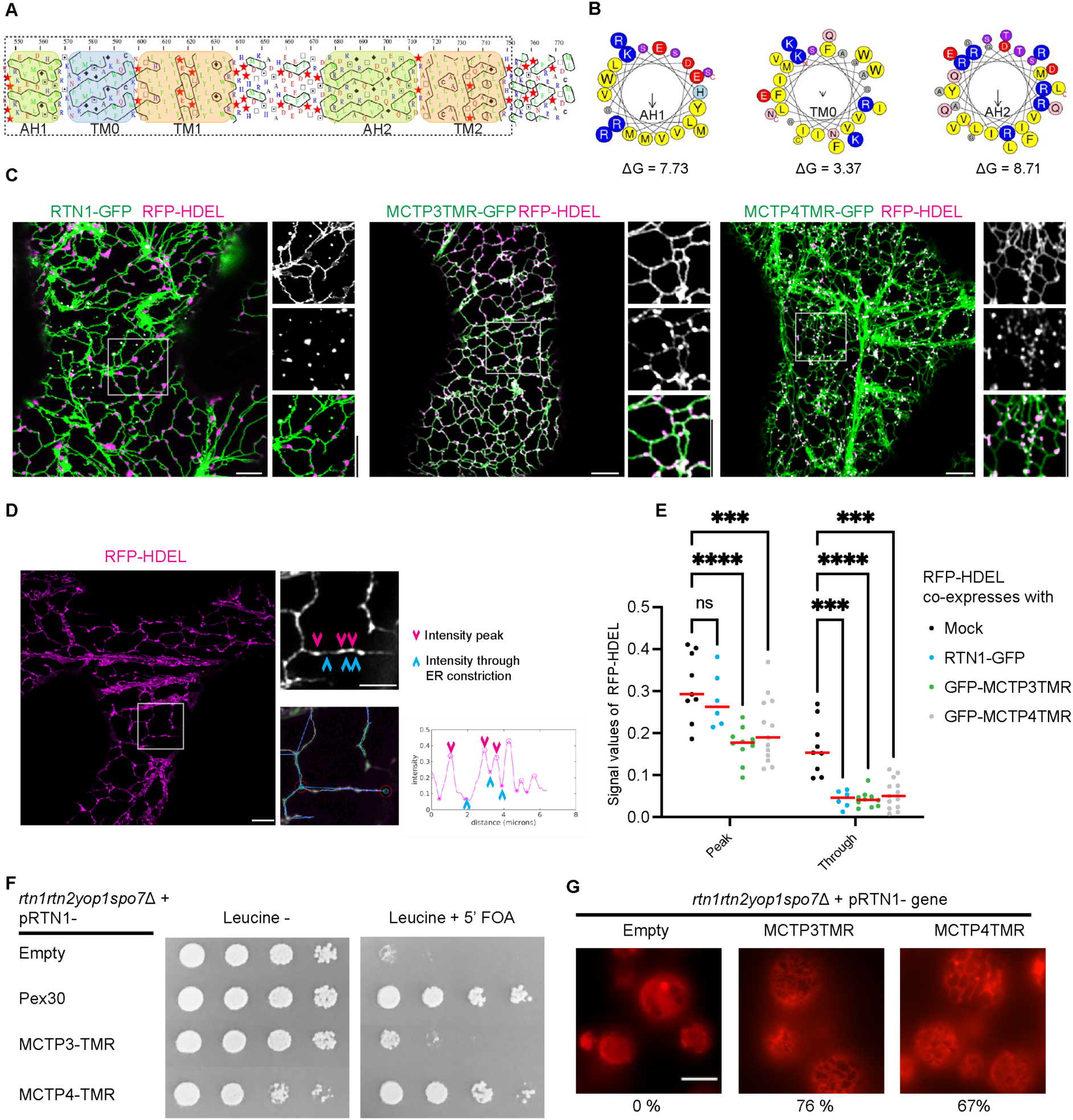
MCTPs are ER shaping proteins. (**A** and **B**) Based on secondary structural analysis, MCTP4 harbors two transmembrane hairpins (TM1 and TM2) and three amphipathic helixes (AH1, TM0 and AH2) at its C-terminus. Altogether, MCTPs transmembrane region (TMR) present similar structural organization and homology to ER-shaping reticulon-homology domain. (**C**) When overexpressed in *N. benthamiana* leaves, MCTP3 and MCTP4 TMR domains cause over-constriction on the ER and loss of luminal space, evidenced by the luminal marker RFP-HDEL being restricted to ER cisternae (as opposed to tubules and cisternae in control conditions). This effect is a hallmark of ER shaping protein, with the ER shaping protein RTN1-GFP shown here as an example. (**D** and **E**) Such constriction effect is further quantitatively reflected by measuring luminal signals along the ER network (*20*). ER is naturally reticulated and has variable lumen spaces (peak and through) along its network. Under conditions when MCTP3TMR, MCTP4TMR, and RTN1 are over-expressed, ER is constricted and causes a reduction of its lumen space, hence a reduction of luminal marker signal which is analyzed through AnalyzER (*20*). n = 9 (HDEL control), n = 6 (RTN1), n = 10 (MCTP3TMR), n = 13 (MCTP4) from 2 technical replicates. Significance was tested using two-way ANOVA test (****, P<0.0001). **(F)** To check if MCTP3/4 TMRs have ER shaping function, we overexpressed them in the conditional lethal *rtn1rtn2yop1spo7*D mutant containing a plasmid with *RTN1* gene and counter selectable marker *URA3*. This mutant is not viable in media containing 5-fluoroorotic acid (5’ FOA) as it kills the cells expressing Ura3. We found that overexpression of Pex30, a reticulon-like ER shaping protein (*39*), and MCTP3/4 TMRs restored the viability of the *rtn1rtn2yop1spo7*D suggesting that MCTP3/4, similar to Pex30, can compensate for the loss of *RTN1* containing plasmid in the 5’ FOA media and have the ER shaping function. **(G)** Images of the rtn1rtn2yop1Δ cells expressing the ER marker ss-RFP-HDEL and MCTP3TMR or MCTP4TMR. Percent cells with normal ER morphology indicated below the images. n = 200 cells. Scale bars: 5 μm (C, D and G).

**Fig. S12.**
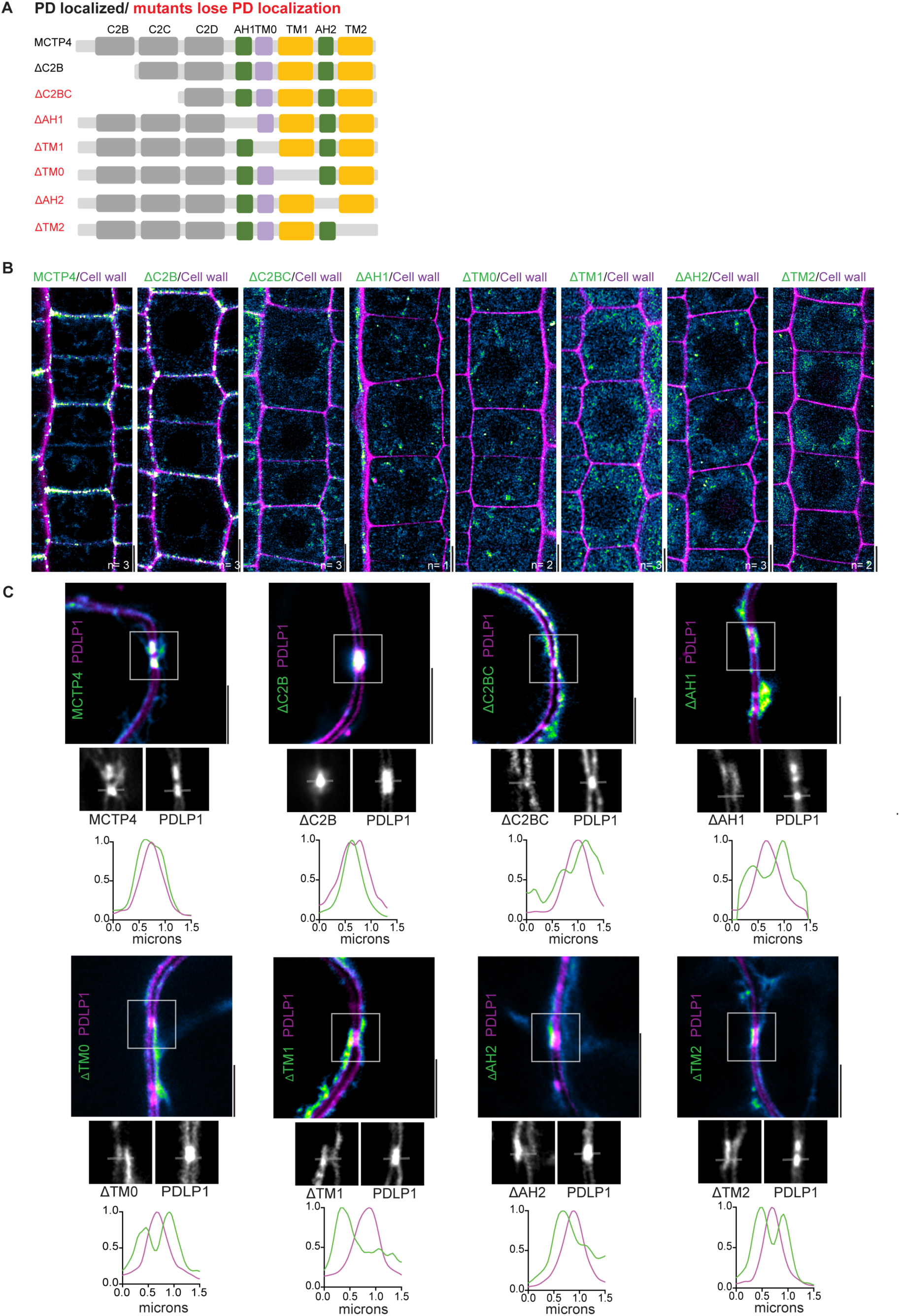
Molecular determinant for MCTP4 plasmodesmata localization. **(A)** Schematic representation of MCTP4 protein domains and the outline of the domain deletion constructs expressed as N-terminal YFP-tagged fusions driven from native MCTP4 promoter in the *mctp3mctp4* mutant (B) or from UBQ10 promoter in *N. benthamiana* leaves together with the plasmodesmata marker *UBQ10:PDLP1mCherry* (plasmodesmata-located protein) (*51*) (C). **(B)** Representative examples of YFP-MCTP4 full-length (FL) and deletions mutant localization (green fire blue) in the primary root epidermal cells in the meristem zone. Cell wall is visualized by propidium iodide (magenta). Number of independent T3 genetic lines observed are indicated below the corresponding images. **(C)** Representative airyscan image of YFP-MCTP4 FL and deletion mutant localization (green fire blue) in the *N. benthamiana* leaf epidermal cells. Plasmodesmata is visualized by the marker PDLP1-mCherry. Localization is consistent across more than 3 technical replicates (B and C). Scale bars, 5 μm (B and C).

**Fig. S13.**
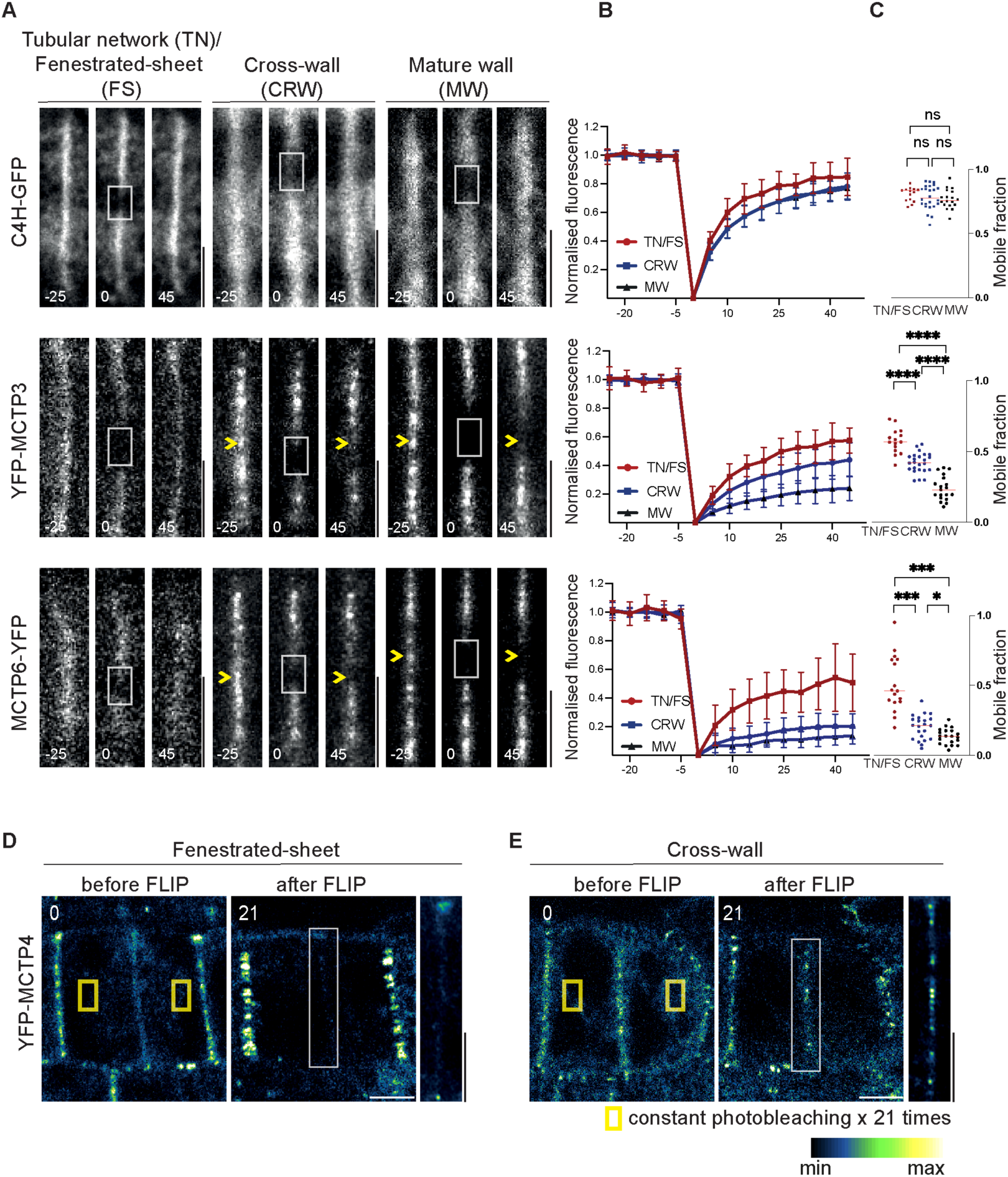
Analysis of MCTPs mobility during and post cytokinesis using photobleaching techniques. **(A)** Cinnamate-4-hydroxylase (C4H), MCTP3 and MCTP6 mobility at tubular network(TN)/fenestrated-sheet (FS), cross-wall (CRW), and mature wall (MW) stages measured by fluorescence recovery after photobleaching (FRAP) in root epidermal meristem cells from Col-0: *35S-C4H-GFP*, *UBQ10-YFP-MCTP3* in *mctp3mctp4* genetically complemented lines and *UBQ10-MCTP6-YFP* in *mctp3mctp4mctp6* complemented seedlings. The outlined regions indicate the photobleaching region at t = 0 s. Yellow arrow indicates MCTP3 and MCTP6 clusters that do not recover fluorescent signal after photobleaching. Time is shown in second. **(B)** quantification of fluorescence (mean ± SD) and **(C)** mobile fraction (bars indicate mean) in n = 16 (C4H), n = 15 (MCTP3), n = 17 (MCTP6) from fenestrated-sheet; n = 24 (C4H), n = 24 (MCTP3), n = 22 (MCTP6) from cross-wall; n =18 (C4H), n = 19 (MCTP3), n = 20 (MCTP6) from mature wall cells across 4 technical replicates. Significance was tested using Kruskal-Wallis one-way ANOVA test. (****, P<0.0001). (**D, E**) Constant photo-bleaching (21 times with 6 sec interval) of YFP-MCTP4 signal in the cortical ER using genetically complemented plants *UBQ10:YFP-MCTP4* expressed in *mctp3mctp4* mutant. Two representative cells: one (D) at fenestrated-sheet when cell plate is about to fuse with the maternal membrane; and (E) cross-wall stage. Yellow square marks bleaching regions and white square marks the cell plate MCTP4 signal at time = 126 s (after 21st bleaching). An airycan image was acquired on the same cell right after the last photobleaching and only cell plate signal was shown here that corresponds to the same region as in t = 126 sec. Note the persistent MCTP4 signal at cross-wall, but not in fenestrated-sheet cells. Scale bars, 5 μm.

**Fig. S14.**
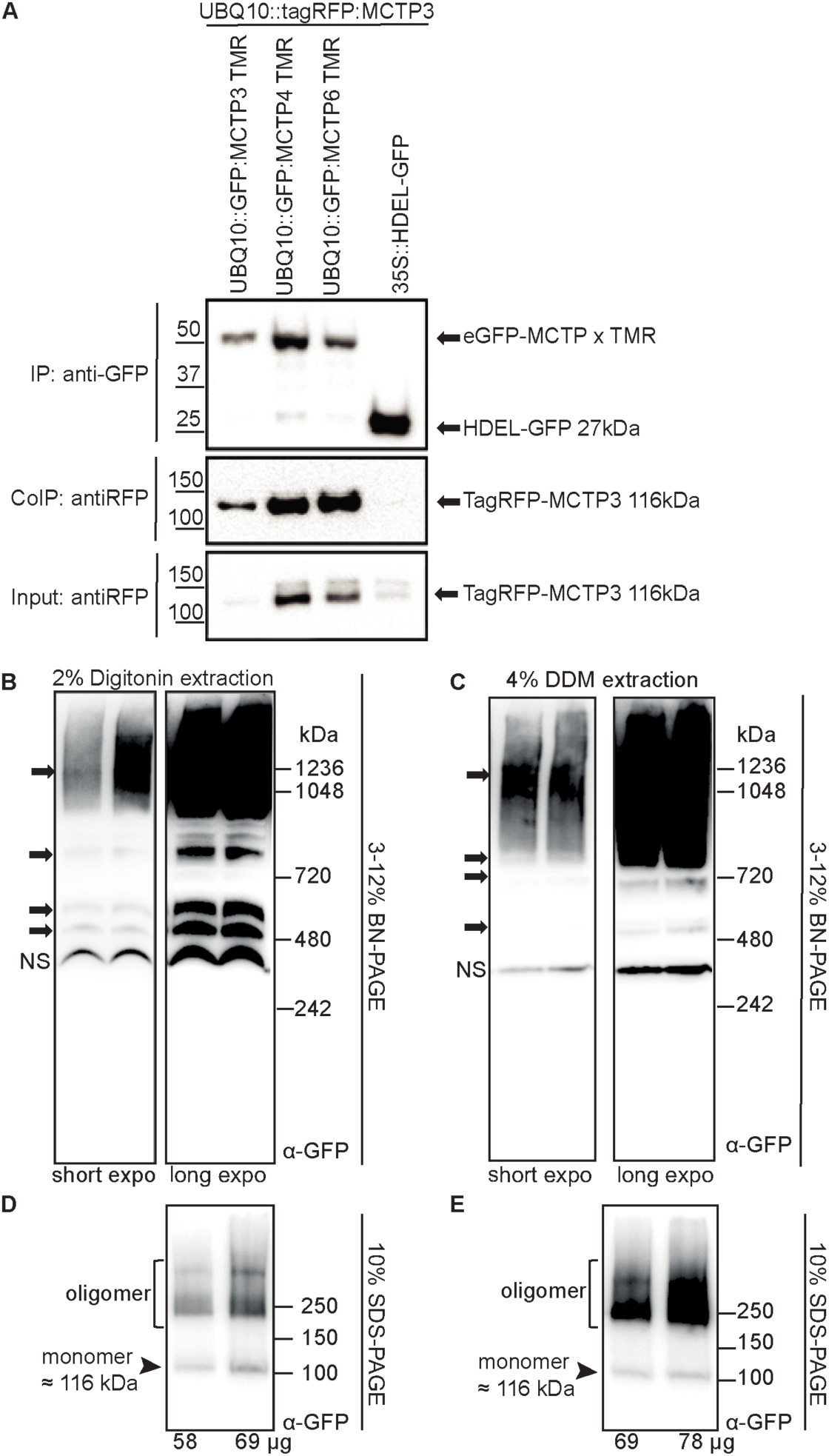
MCTPs form high-order oligomers. **(A)** Co-immunoprecipitation between pray protein tagRFP-MCTP3 and baits GFP-MCTP3 TMR, GFP-MCTP4 TMR, GFP*-*MCTP6 TMR and GFP-HDEL transitory expressed in *N. benthamiana* leaves. Representative results from two technical replicates. **(B-E)** *UBQ10:YFP-MCTP4* expressed in *mctp3mctp4* complemented Arabidopsis seedlings, extracted by digitonin or DDM, oligomerize and form high molecular weight (MW) complexes in vivo, as shown by 3-12 % BN-PAGE (B-C) and 10% SDS-PAGE (D-E) western blot analysis (using anti-GFP antibody). Black arrows (B-C) point to the MCTP4 high MW complexes in both short (51 s) and long exposed (157 s) blots, black arrowheads (D-E) points to the monomer MCTP4 that is denatured by the SDS. NS indicates non-specific band. Total protein quantities run in the gel are indicated at the bottom. Representative results from more than three technical replicates.

**Fig. S15.**
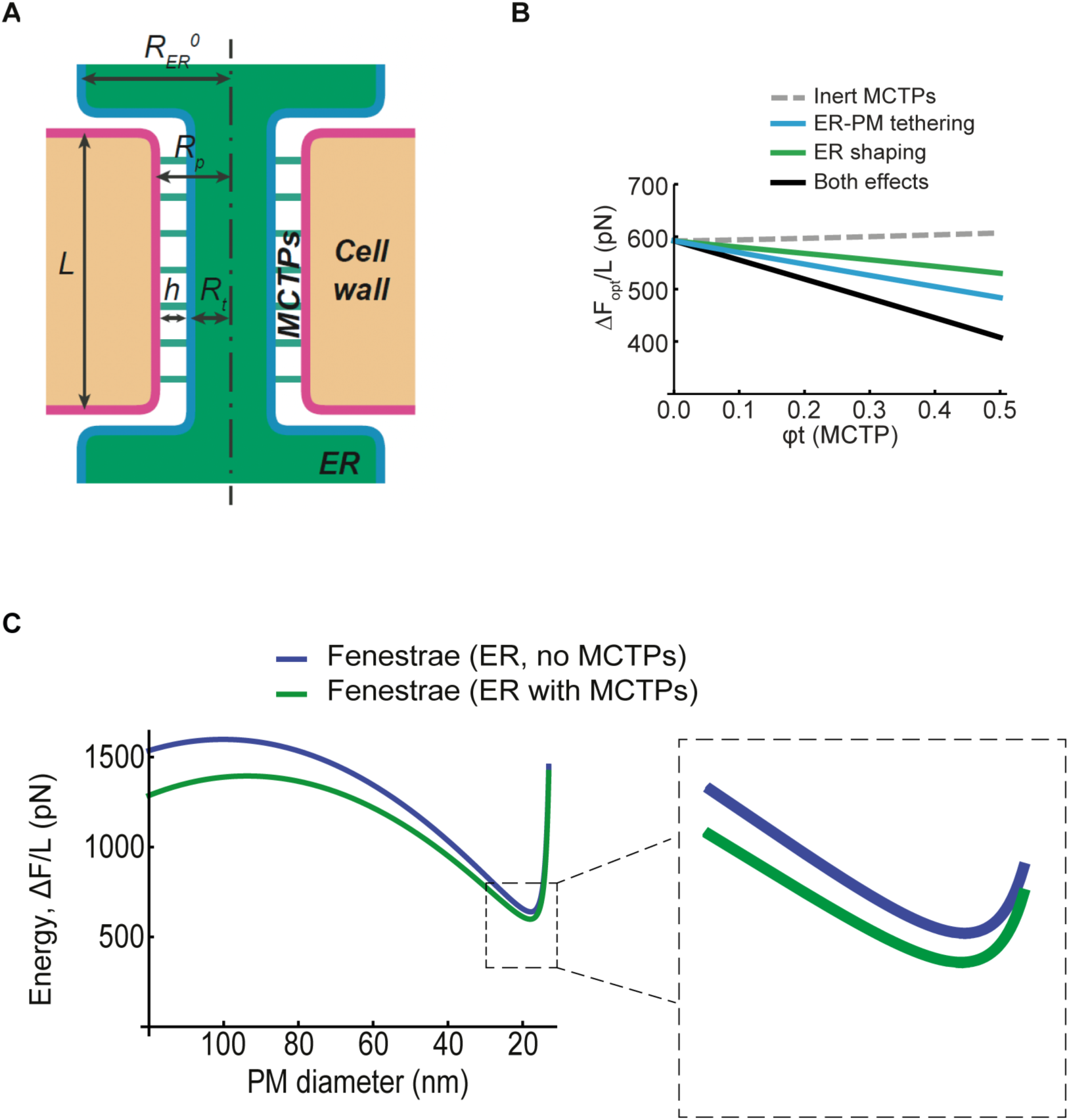
Semi-quantitative physical model of nascent plasmodesmata morphology and energetics. **(A)** Schematic model of one plasmodesma. *L*, longitudinal length; 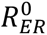 and *R*_*t*_, ER radius outside and inside forming plasmodesmal bridge/fenestrae. *R*_*p*_, radius; *h*, distance between the cell plate membrane (future PM) and the ER inside plasmodesmata bridged by MCTP proteins (green lines connecting the two membranes). **(B)** Physical model showing that MCTP enrichment (ϕ_t_=fraction of MCTP covering the ER surface) stabilizes nascent plasmodesmata by reducing the free energy of the system, depending on the tethering and ER-shaping functions. **(C)** The presence of ER works against fenestrae closure by locally creating a stable structure (metastable) of about 20 nm in diameter (ΔF, free energy of the structure. L, longitudinal length of forming plasmodesmata). The presence of MCTPs (green line) decreases the free energy of the metastable state (dip in the curve) compared to ER with no MCTPs (blue line). Lower energy means that the free energy barrier is larger (the jump that needs to be overcome is larger), meaning that the system is more stable with MCTPs than without. (Inset: zoom-in of the free energy curves).

**Fig. S16.**
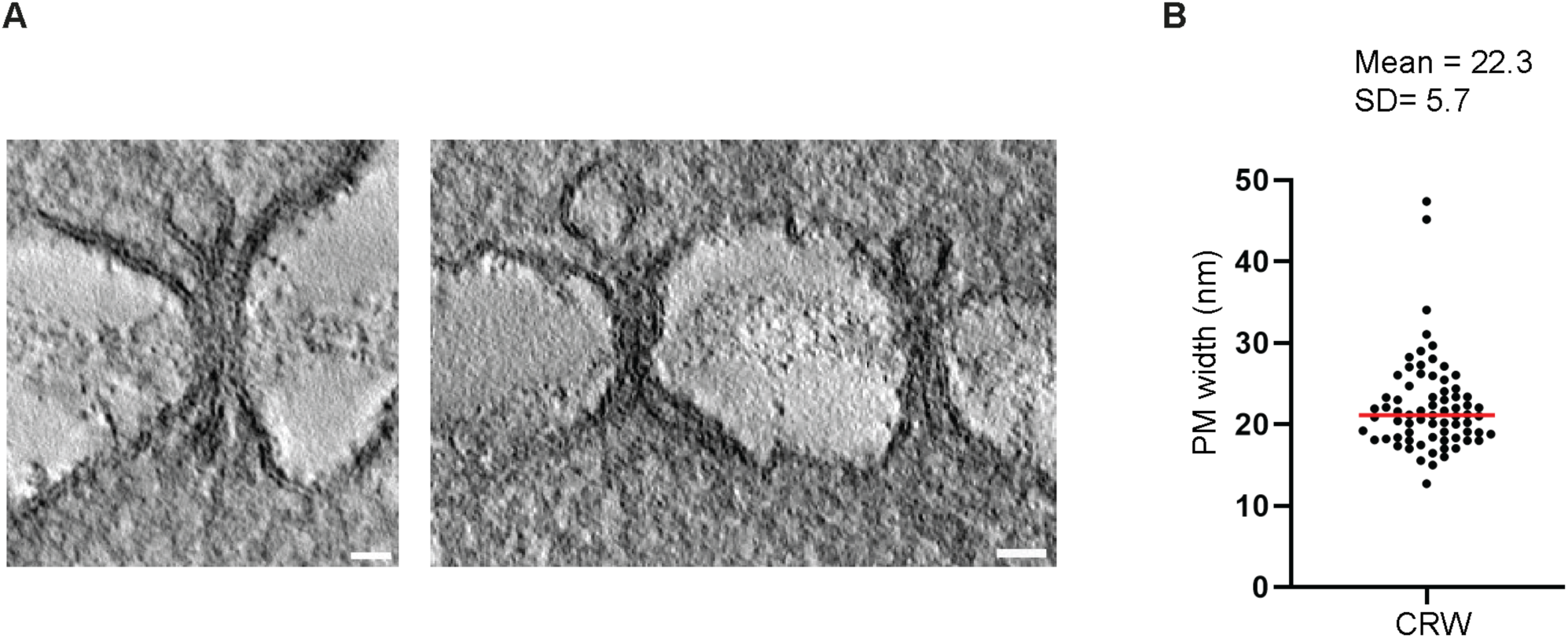
Measurement of nascent plasmodesmata diameter in *mctp3mctp4mctp6* mutant. (**A**) Reconstructed scanning transmission electron tomography of plasmodesmata in the *mctp3mctp4mctp6* mutant root meristem endodermis cells at the cross-wall stage. (**B**) Diameter of stabilized fenestrae at CRW stage, n = 72, extracted from electron tomography acquisition as exemplified in (A). Scale bars, 50 nm.

**Table S1.** Parameters used for the physical modeling.

| Parameter | Value | Reference |
| --- | --- | --- |
| Radius of the PM fenestra, $R_p$ | Free parameter | |
| Radius of the ER tubules, $R_t$ | Free parameter | |
| Tube MCTP area fraction, $\phi_t$ | Free parameter | |
| Plasmodesmata length, $L$ | 150 nm | (18) |
| Cell wall constricting force, $f_{constr}$ | 2–10 pN/nm<br>(10 pN/nm unless otherwise stated) | (52) |
| Osmotic pressure, $\Delta p$ | 0.01–0.2 pN/nm<sup>2</sup> (0.2 pN/nm<sup>2</sup> unless otherwise stated) | (53) |
| Bending rigidity, $\kappa$ | 80 pN nm | (54) |
| Plasma membrane tension, $\sigma_{PM}$ | 0.1 pN/nm | (55) |
| MCTP cross-sectional surface area, $a_{MCTP}$ | 10 nm<sup>2</sup> | *Estimation based on molecular sequence/structure* |
| Bulk MCTP area fraction, $\phi_b$ | <0.001 (endogenous levels; higher for overexpressing conditions) | *Estimation based on standard ER protein densities* |
| ER tube spontaneous curvature, $J_{s,0}$ | 0.036 nm<sup>-1</sup> (obtained using) | (56) |
| ER membrane tension, $\sigma_{ER}$ | 0.012 pN/nm | (57) |
| MCTP effective spontaneous curvature, $\zeta_{MCTP}$ | 0–0.5 nm<sup>-1</sup><br>(0.5 nm<sup>-1</sup> unless otherwise stated) | *Typical values for powerful curvature generating proteins*<br>(43) |
| MCTP PM binding energy, $\epsilon_a$ | 10 pN nm | *Typical values for powerful protein-lipid binding energies* |
| MCTP tethering reach, $h_0$ | 10 nm | *Estimation based on molecular sequence/structure* |
| Degree of membranes hydration, $P_0$ | 200 pN/nm<sup>2</sup> | (49) |
| Hydration length, $\xi_h$ | 0.25 nm | (49) |
| Monolayer thickness, $\delta$ | 2 nm | (58) |

**Table S2.**
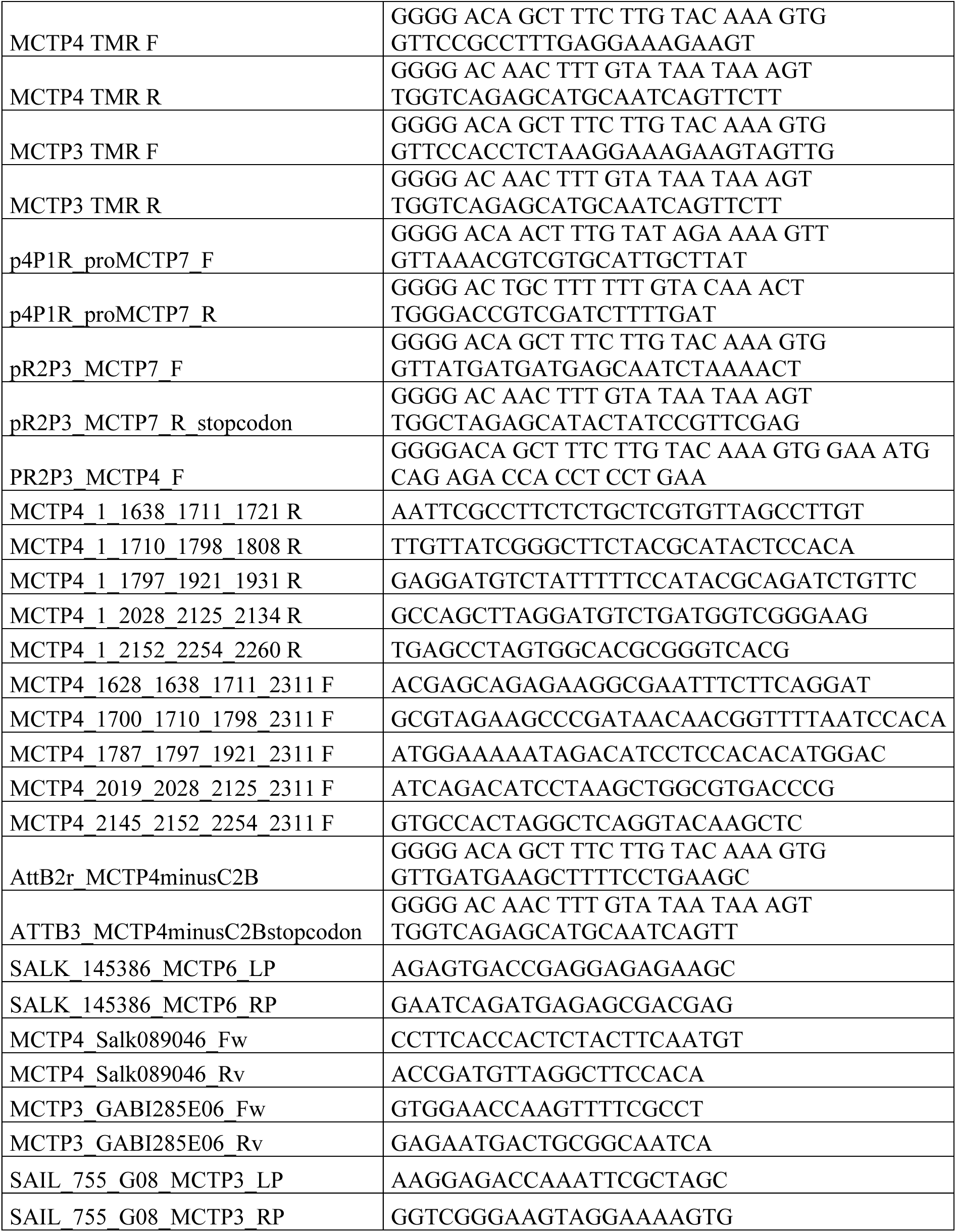
List of primers.

